# SUMOylation of Dorsal attenuates Toll/NF-κB signalling

**DOI:** 10.1101/2021.09.07.459254

**Authors:** Sushmitha Hegde, Ashley Sreejan, Chetan J Gadgil, Girish S Ratnaparkhi

**Affiliations:** Biology, Indian Institute of Science Education & Research, Pune 411008, INDIA; Chemical Engineering and Process Development Division, CSIR - National Chemical Laboratory, Pune 411008, INDIA; CSIR-Institute of Genomics and Integrative Biology, New Delhi 110020, INDIA

**Keywords:** *Drosophila*, Haploinsufficiency, innate immunity, SUMO, transcription

## Abstract

In *Drosophila*, Toll/NF-κB signalling plays key roles in both animal development and in host defence. The activation, intensity and kinetics of Toll signalling is regulated by post-translational modifications such as phosphorylation, SUMOylation or ubiquitination that target multiple proteins in the Toll/NF-κB cascade.

Here, we have generated a CRISPR-Cas9 edited Dorsal (DL) variant that is SUMO conjugation resistant (SCR). Intriguingly, embryos laid by *dl^SCR^* mothers overcome *dl* haploinsufficiency and complete the developmental program. This ability appears to be a result of higher transcriptional activation by DL^SCR^. In contrast, SUMOylation dampens DL transcriptional activation, ultimately conferring robustness to the dorso-ventral program. In the larval immune response, *dl^SCR^* animals show increase in crystal cell numbers, stronger activation of humoral defence genes, high *cactus* levels and cytoplasmic stabilization of DL:Cactus complexes. A mathematical model that evaluates the contribution of the small fraction of SUMOylated DL (<5%) suggests that it acts to block transcriptional activation, driven primarily by DL that is not SUMO conjugated.

Our findings define SUMO conjugation as an important regulator of the Toll signalling cascade, in both development and in host defense. Our results broadly indicate that SUMO attenuates DL at the level of transcriptional activation. Further, we hypothesize that SUMO conjugation of DL may be part of a Ubc9 dependant feedback circuit that restrains Toll/NF-κB signalling.

## Introduction

Toll-like receptor (TLR) signalling is a highly conserved, ancient response to combat pathogenic attacks in multicellular eukaryotes (Kopp and Ghosh 1995; Medzhitov *et al*. 1997; Zhang and Ghosh 2001; Janeway and Medzhitov 2002). In *Drosophila*, in addition to its role in regulating the host response to infection, the Toll/Dorsal pathway has been co-opted to orchestrate early development, laying down the foundations for the dorso-ventral (DV) body plan (reviewed by (Steward and Govind 1993; Morisato and Anderson 1995; Belvin and Anderson 1996; Rusch and Levine 1996; Stathopoulos and Levine 2002; Valanne *et al*. 2011). The chief effector in DV development, the NF-κB transcription factor Dorsal (DL) is held inactive in the cytoplasm by the IκB ortholog Cactus (Cact) (Steward 1987; Roth *et al*. 1989, 1991; Rushlow *et al*. 1989; Geisler *et al*. 1992; Govind *et al*. 1993; Whalen and Steward 1993). The binding of the ligand Spaetzle (Spz) to the Toll receptor sets in motion a kinase cascade, leading to the degradation of Cact and nuclear translocation of DL (Bergmann *et al*. 1996). In the early embryo, the partitioning of maternal DL in the ventral nuclei creates a DV gradient dictated by the strength of activation of the Toll receptor (Steward *et al*. 1988; Roth *et al*. 1989).

Asymmetric Toll signalling initiates the DL gene regulatory network, activating 50-70 target genes to specify the presumptive germ layers of the fly (Hong *et al*. 2008). Peak levels of DL activate *twist (twi)* and *snail (sna)* in the ventral most regions, specifying the mesoderm (Ferguson and Anderson 1991; Ray *et al*. 1991). Lower levels of DL in the lateral regions turn on genes like *rhomboid (rho), brinker (brk)* and *short gastrulation (sog)*, giving rise to the neuro-ectoderm. DL also acts as a repressor of ectodermal genes, like *zerknullt (zen)* and *decapentaplegic (dpp)*, in the presumptive mesoderm and neuroectoderm (Kosman *et al*. 1991, 1992; Ip *et al*. 1992; Araujo and Bier 2000).

The Toll signalling arm is also deployed later in the *Drosophila* life-cycle to ward off fungal and gram-positive bacterial insults, triggering the production of a battery of defense genes, which include antimicrobial peptides (AMPs) as part of the systemic humoral immune response (Lemaitre *et al*. 1996; Rutschmann *et al*. 2002; Ferrandon *et al*. 2007). Though DL-related Immunity Factor (Dif) is the primary immune NF-κB effector in the Toll pathway, DL acts in concert with Dif to aid in host defense (Lemaitre *et al*. 1995; Anderson 2000). Toll/NF-κB signalling is subject to regulation by post-translational modifiers (PTMs) (Karin and Ben-Neriah 2000; Zhou *et al*. 2005). The best characterized example is the phosphorylation of Cact in response to activation of the receptor, which leads to degradation of Cact and release of DL for its journey to the nucleus (Belvin *et al*. 1995; Liu *et al*. 1997). DL itself is subject to regulation by phosphorylation, where the modification contributes to its graded nuclear import in the embryo (Gillespie and Wasserman 1994; Drier *et al*. 1999).

SUMO resembles ubiquitin in its three-dimensional structure but utilizes a related but unique set of conjugating enzymes to attach to target proteins at a lysine residue, most often part of a ψ-K-X-E/D consensus motif (Johnson 2004; Hay 2005; Varejão *et al*. 2020). The covalent conjugation of SUMO to targets confers novel properties, in terms of altered localization, stability, or activity. The modified protein can further be recognized by a cognate partner via a SUMO-interaction motif (SIM), thus changing its interaction potential (Gareau and Lima 2010). SUMO regulates a plethora of cellular processes and is upregulated in response to protein-damaging stresses like heat shock, osmotic stress, proteasomal inhibition, immune stress, etc (Enserink 2015; Hegde *et al*. 2020). This increase in global SUMOylation, termed the SUMO-stress response, is an essential cyto-protective adaptation (Ryu *et al*. 2020).

DL has been shown to be a SUMO target, based on experiments conducted in *Drosophila* S2 cells (Bhaskar *et al*. 2000, 2002). Studies in larvae have also emphasized the interplay of SUMO and the Toll pathway in modulating host defense. Mutations in the SUMO E2 ligase Ubc9, encoded by *lwr* in *Drosophila*, lead to the over-proliferation of hemocytes. Introducing mutations in the *dl* and *Dif* loci in a *lwr* mutant background restores the wild type blood cell population, providing evidence for the intersection of Toll signalling with the SUMO conjugation machinery (Huang *et al*. 2005; Chiu *et al*. 2005). In the embryo, mass spectrometric studies suggest that DL is SUMO conjugated (Nie *et al*. 2009). However, roles for DL SUMOylation in the animal have not been studied.

Armed with these pieces of evidence, we sought to delineate the function of SUMO conjugation of DL in *Drosophila*. In our study, we address the effect of SUMO conjugation on DL function by employing a CRISPR-Cas9 based strategy, which allows precise editing of the DL locus, replacing the 382^nd^ lysine with a charge-preserving arginine. This *dl^K382R^* animal is then subsequently evaluated for its effect on early development and host-defense, both of which represent critical spatiotemporal domains for Toll/DL signaling. Our studies uncover roles for SUMO conjugation of DL in supporting the robustness of embryonic DV patterning. Further, we find that DL SUMOylation has roles in both the cellular and humoral response in the larvae. In these two distinct signaling contexts, a common mechanism that emerges is the role of DL SUMO conjugation in negatively regulating Toll signaling by specifically attenuating DL mediated transcriptional activation.

## Results

### Generation of a genome-edited *dl^K382R^* mutant

In recent years, the advent of CRISPR-Cas9 genome editing technology has allowed the generation of point mutations in a straightforward and site-directed manner (Bassett *et al*. 2014; Gratz *et al*. 2014; Bier *et al*. 2018). DL has a single, well characterized, and validated SUMO conjugation site at K382 (Fig. 1A) (Bhaskar *et al*. 2002). The K382R mutation abolishes SUMO conjugation, generating a SUMO-conjugation-resistant (SCR) variant of DL. We employed the following protocol (Fig 1A, B; Materials and Methods) to generate a *dl^K382R^* mutation (Fig 1C, D). A single guide RNA (sgRNA) targeting the *dl* locus - with no predicted off-target cleavage sites and designed to induce a double-strand break at *dl^K382^*, in the presence of Cas9, was cloned into the pBFv-U6.2 plasmid. A 100 bp-long ssODN (Fig. 1A) harbouring the K382R mutation was supplied as the repair template and co-injected along with the sgRNA plasmid in embryos expressing Cas9 in the *vasa* domain. Details of the cloning of the gRNA are described in the methods section. The 450 flies (F0) that emerged from the injected embryos were crossed to a second chromosome *w*^-^;*Tft/CyO* balancer (Fig. 1B). Three animals from each vial, for each of the 400 lines, were crossed again to *w*^-^;*Tft/CyO*, to generate stable, putative, *dl^K382R^/*Cyo lines. Of these, 200 homozygous, putative transformants were screened for insertion of the ssODN by PCR amplification of the genomic locus and subsequent restriction digestion of the PCR product to identify the mutated nucleotide sequence (Fig. 1C). The screening strategy incorporated a site for the restriction enzyme BstBI in the ssODN, validating the successful incorporation of the mutation in the genome. Based on restriction digestion patterns, ∼2% of lines (4 out of 200), harboured the mutation and we validated four of these (26.1, 110.1, 242.1, 266.1) by sequencing (Fig. 1C). Representative sequencing data is shown in Fig. 1D. A few lines containing wild-type sequences were also retained and one of these (72.1) was defined as a ‘CRISPR-control’, *dl*, at par with the wild type animal. All *dl^K382R^* and *dl* lines were homozygous viable with comparable *dl* transcript levels across the embryonic (Fig. S1A), larval (Fig. S1B), and adult stages (Fig. S1C).

**Fig. 1.**
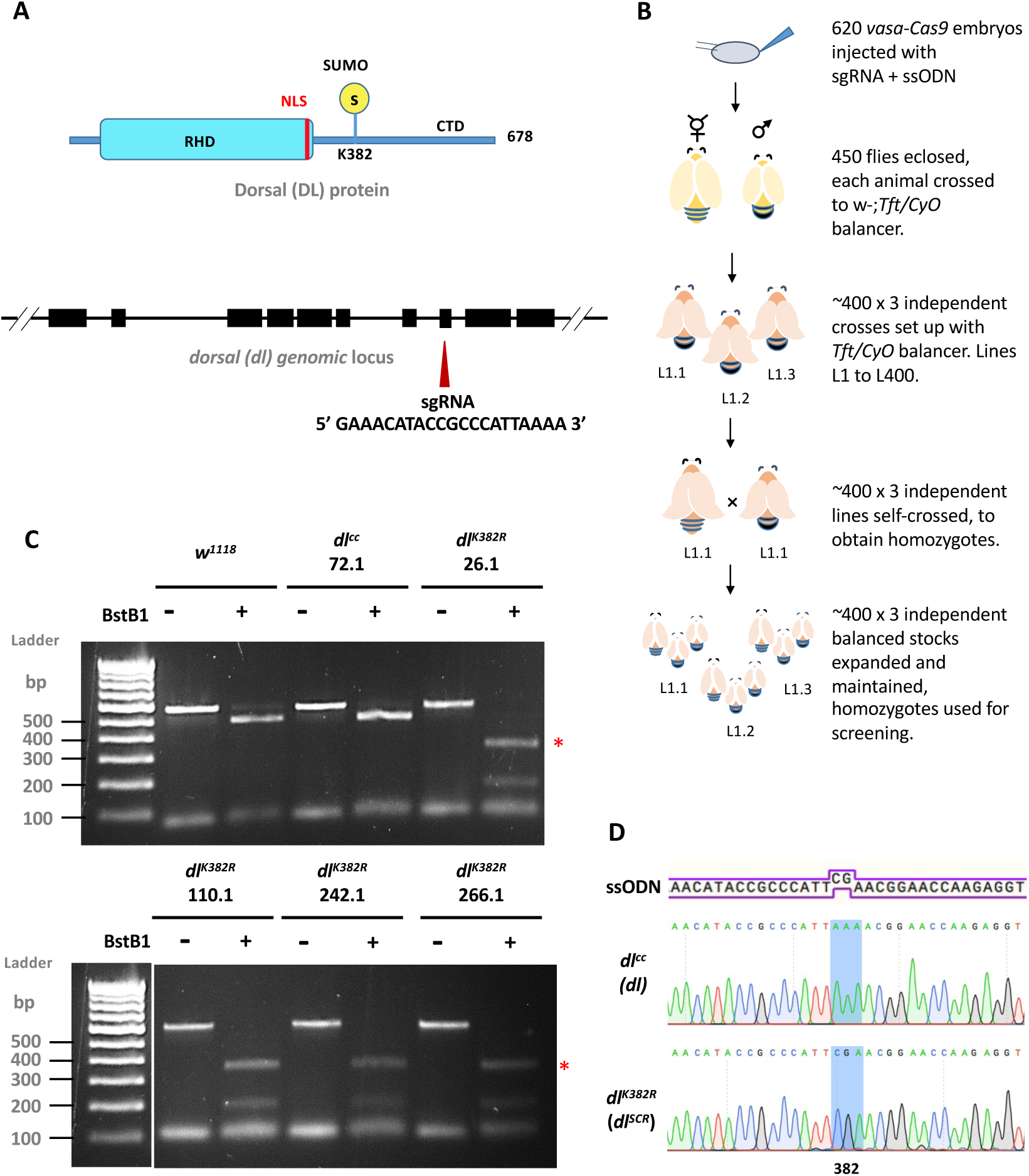
Creating the *dl^K382R^* mutant using the CRISPR-Cas9 system. A schematic of the DL protein (in blue) and gene locus (in black) is presented in panel (A). DL is SUMOylated at K382, part of the consensus motif I**K**TE. A 20 bp *sgRNA* was designed to create a double strand break in the vicinity of *dl^K382^,* downstream of the genomic region coding for DL Rel-homology domain (RHD) and nuclear localization sequence (NLS), in exon 8. The detailed crossing scheme for the generation of the *dl^K382R^* allele after injection of the gRNA plasmid and ssODN is outlined in (B). Homozygous flies obtained were screened by genomic PCR and digestion with the BstB1 enzyme, which recognizes the engineered site of mutation, TTCGAA (C). Four independent lines - 26.1, 110.1, 242.1 and 266.1 showed a distinct digest of the PCR product (indicated by red asterisks), while line 72.1 served as a control. The presence of the mutation was confirmed through sequencing, and the K382R mutation, encoded by the codon CGA, is highlighted in blue, in the representative chromatogram (D).

The *dl^K382R^* genome-edited lines were also used in a trans-allelic combination (e.g 26.1/110.1) to negate off-target effects. Here onwards, *dl^K382R^* is referred to *dl^SCR^*, a line where DL is resistant to SUMO conjugation.

### Early development proceeds normally in *dl^SCR^* embryos

The initial hours of embryonic development are driven by the deposition of maternal gene products (Anderson and Nüsslein-Volhard 1984; St Johnston and Nüsslein-Volhard 1992; Horner and Wolfner 2008; Tadros and Lipshitz 2009; Stein and Stevens 2014; Laver *et al*. 2015). DL is deposited maternally and functions as a master regulator in specifying the DV axis (Santamaria and Nüsslein-Volhard 1983; Roth *et al*. 1989; Ratnaparkhi and Courey 2015). DL is also among the ∼140 maternal proteins identified as substrates for SUMO conjugation in the 0-3 hour embryo, using mass-spectrometry (Nie *et al*. 2009), and the *dl^SCR^* fly line is ideal to uncover roles for SUMO conjugation of DL. The DL nucleo-cytoplasmic gradient, which could be influenced by SUMO conjugation, appears to be normal (Fig. 2A), in eggs laid by homozygous *dl^SCR^* mothers. Also, cuticle patterns, which are a sensitive readout for aberrations in both maternal and zygotic stages of development were normal in *dl^SCR^* embryos (Fig. 2A). We observed that embryos deposited by homozygous *dl^SCR^* mothers show no discernible changes in embryonic viability (Fig. 2B), when compared to controls. In addition, we measured levels of transcripts of *dl* and its primary ventral targets *twist, snail* and *zen* by quantitative real-time polymerase chain reaction (qRT-PCR). We failed to detect significant changes in these transcripts, except *zen*, in the *dl^SCR^* mutants, in comparison to the control (Fig. 2C). Taken together, these results demonstrate that DL SUMOylation is either dispensable or that the effect of the *dl^SCR^* mutation is compensated for by unknown mechanisms in the developing embryo.

**Fig. 2.**
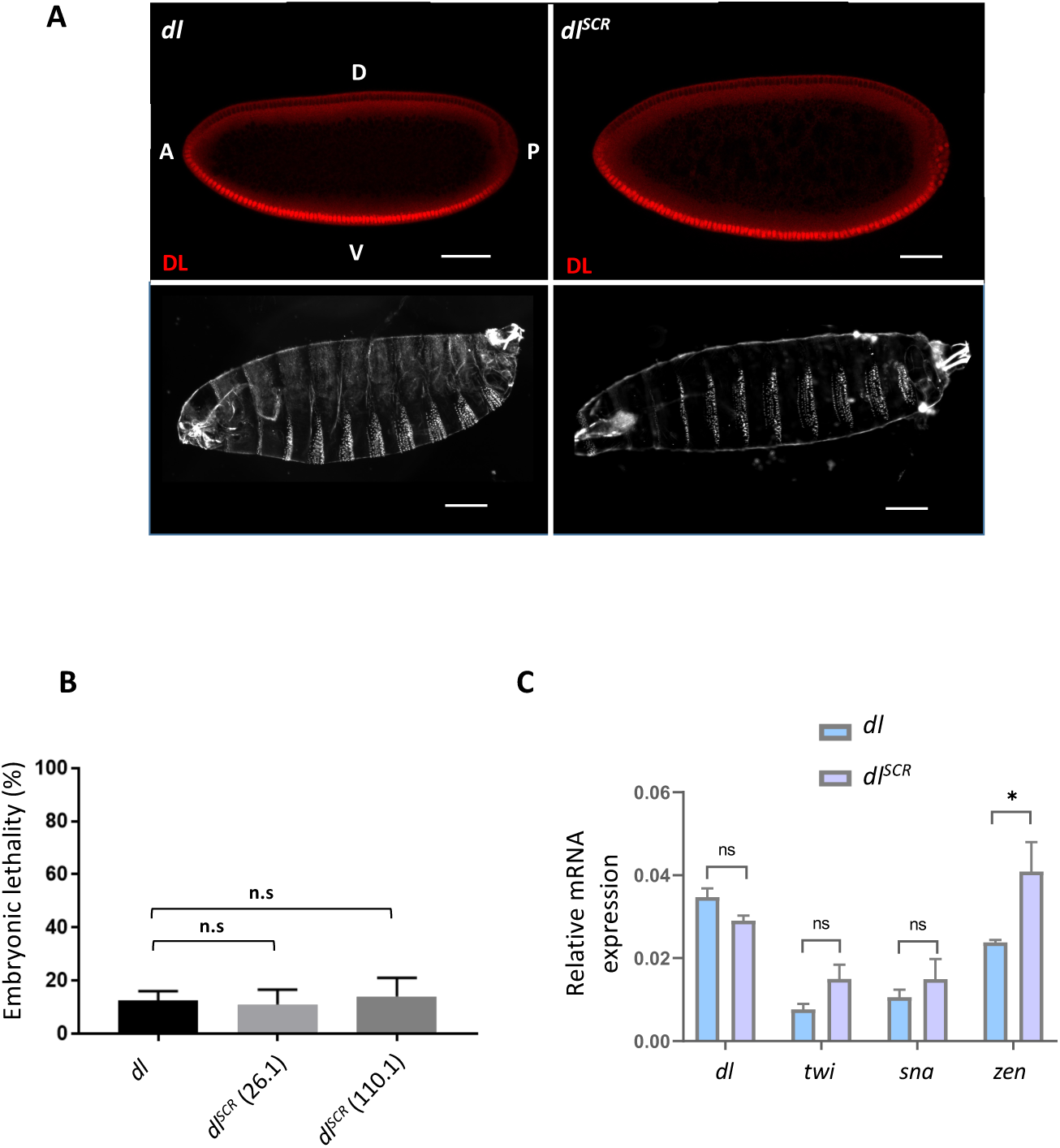
SUMO conjugation is dispensable for embryonic development. Syncytial blastoderm embryos were stained for DL and localization in the nuclei observed (A, top panel). Embryos are oriented dorsal-side up and anterior-side on the left. Cuticle preparations (A, bottom panel) indicate regular arrangement of denticle bands. Percentage of unhatched embryos is plotted as embryonic lethality for control and two of the mutant lines, 26.1 and 110.1. Genotype of mated mothers is listed on the X-axis (B). N=3, ordinary one-way ANOVA, (ns) *P* >0.05. (C) represents qRT-PCR analysis of *dl* transcripts and DL target genes *twi*, *sna* and *zen* for embryos from mated females of the genotypes *dl* and *dl^SCR^*. N=3, Two-way ANOVA, (ns) *P* >0.05, (*) *P* <0.05.

### Haploinsufficiency of *dl* is rescued in ***dl^SCR^*** embryos

Since SUMO is essential to adapt to a multitude of cellular stresses, we reasoned that a requirement for SUMOylation of DL would only be apparent under conditions of stress. Temperature stress is well documented to induce an increase in global SUMOylation levels (Tempé *et al*. 2008). Moreover, SUMOylation of DL was observed in heat-shocked embryos (Nie *et al*. 2009). The embryonic lethality is comparable at 29 °C, for *dl* and *dl^SCR^* (Fig S2A). A well-known allelic combination that disrupts the DV developmental program is *dl* haploinsufficiency at 29 °C, where ∼50% of embryos fail to hatch into larvae (Nüsslein-Volhard 1979; Nüsslein-Volhard *et al*. 1980; Simpson 1983). To generate this sensitized background, we crossed *dl* animals to a deficiency line, J4, designated as Df henceforth, where the *dl* and *dif* genomic loci are deleted (Meng *et al*. 1999). As expected, only ∼50% of eggs laid by *dl/Df* lines could hatch at 29 °C. Surprisingly, the embryonic lethality of *dl^SCR^/Df* was significantly lower, at 15% in comparison to the 55% lethality of *dl/Df* embryos (Fig. 3B). This result was consistent across two of the *dl^SCR^* lines tested, 26.1 and 110.1. Similar results were obtained in combination with *dl^1^*, another null allele of *dl* (Fig. S2B). At 25 °C, the embryonic lethality for both *dl^SCR^/dl^1^* and *dl/dl^1^* were comparable (Fig. 3A), indicating that a reduction of *dl* gene dosage at 25 °C is sufficiently well-tolerated, unlike at 29 °C.

**Fig. 3.**
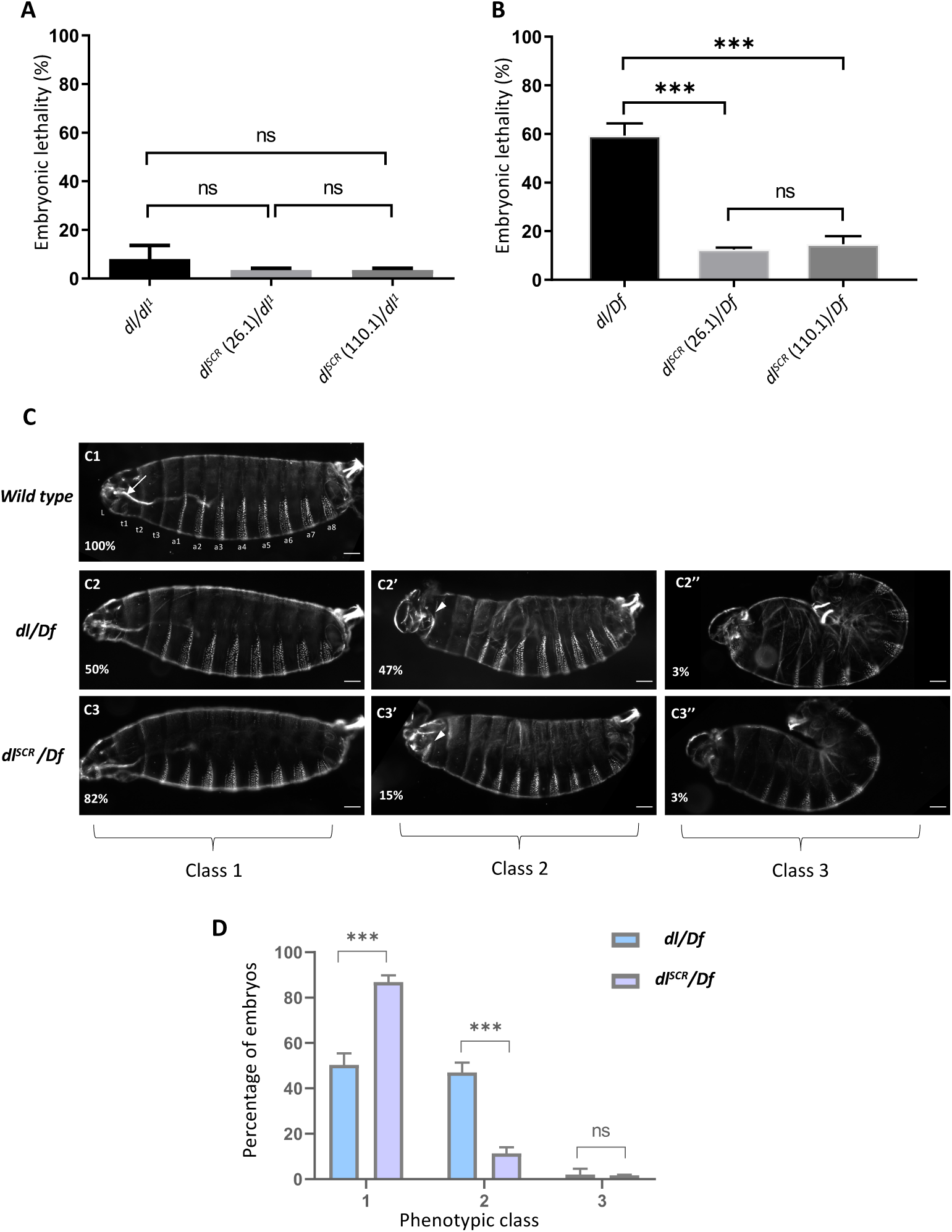
*dl^SCR^ is* haplo-sufficient. Progeny of mothers of the indicated genotype, with one functional copy of DL, were scored for viability, 48 hours after egg lay, at 25 °C (A) and 29 °C (B). N=3, mean ± SEM, Ordinary one-way ANOVA, (ns) *P* >0.05, (***) *P* < 0.001. Cuticle preparations of progeny of the maternal genotypes *dl/Df* and *dl^SCR^/Df,* visualized under a dark-field microscope yielded three major ranges of phenotypes, classified as class 1 (intact head region/ mouth hook; ventral denticle bands and filzkorper normal), class 2 (defective head structure; denticle bands and filzkorper intact) and class 3 (twisted embryos; defective head structures and filzkorper) (C). Cuticles are oriented dorsal-side up and anterior-side on the left. >100 embryos were scored in each replicate, and the percentage of each phenotypic class is plotted for *dl/Df* and and *dl^SCR^/Df* (D). N=3, mean ± SEM, Two-way ANOVA, (ns) *P* >0.05, (***) *P* < 0.001.

To determine if the lethality due to haplo-insufficiency was the consequence of an underlying deficit in DL-mediated patterning, we turned our attention to the cuticles of first instar larvae. The wild-type larval cuticle displays a regular arrangement of ventral denticle bands with intact anterior head structures and terminal Filzkorper (Wieschaus and Nüsslein-Volhard 2016). Eight distinct abdominal segments (a1-a8), three thoracic segments (t1-t3), and an anterior labial segment (L) are demarcated in the wild-type cuticles, hereafter designated as phenotypic Class I (Fig. 3-C1). 50% of *dl/Df* embryos, in contrast to 87% of *dl^SCR^/Df* embryos present as Class I (Fig. 3-C2, C3, D), concurrent with the percentage of embryos that hatch. 47% of *dl/Df* embryos show a reduced head skeleton, and missing labial and T1 segments while the abdominal denticle bands and Filzkorper remained intact (Fig. 3-C2’, C3’). Embryos displaying this phenotype, defined as Class II, were drastically reduced in *dl^SCR^/Df*, to 11%. A small fraction of embryos (2% of *dl/Df* and 1.7% of *dl^SCR^/Df*) showed a more severe phenotype (Fig. 3-C2’’, C3’’). They present with a twisted morphology, defective head structures, and misplaced Filzkorper referred to as Class III, reminiscent of weakly dorsalized D3 embryos. The rescue of embryonic lethality appeared to be a direct result of Class II embryos transiting to normal, Class I embryos in the presence of the *dl^SCR^* allele. Our findings suggest that the *dl^SCR^* allele alleviates temperature-dependent haploinsufficiency, rescuing developmental patterning.

### DL^SCR^ supports the developmental program under haploinsufficient conditions

In the DV axis, DL activates and represses 50-70 genes in the syncytial embryo, influencing gene networks that later define mesodermal, neuroectodermal and ectodermal cell fates (Maggert *et al*. 1995; Markstein *et al*. 2002; Stathopoulos and Levine 2002; Cowden and Levine 2003; Papatsenko and Levine 2005, 2008; Zeitlinger *et al*. 2007; Ratnaparkhi and Courey 2015). These gene networks are a result of a number of factors that include the DL gradient (Fig. 4, A1), variation in strength and number of DL binding sites on promoters of target genes, DL/Twist synergy and activation of positive and negative regulators (Reeves and Stathopoulos 2009). A majority of genes regulated by DL are activated, e.g. *twist(twi), snail (sna), rhomboid (rho), brinker (brk), short gastrulation (sog),* while a small number, such as *decapentaplegic* (*dpp)* and *zerknullt (zen)* are transcriptionally repressed. A number of genes are activated/repressed in spatiotemporal patterns that are very well characterized (Moussian and Roth 2005). *twi* is one of the earliest targets of DL to be activated in the ventral region in the wild type embryo (Fig 4, B1) (Jiang *et al*. 1991; Kosman *et al*. 1992). In haploinsufficient embryos, the DL gradient has been measured to be shallower and broader (Liberman *et al*. 2009; Reeves *et al*. 2012; Ambrosi *et al*. 2014; Carrell *et al*. 2017; Al Asafen *et al*. 2020). In agreement, we find weaker staining of DL (Fig. 4A’, A’’) in haploinsufficient embryos, at 29 °C. A large fraction (∼40%) of embryos laid by *dl/Df* mothers deviate from normal reduction or complete disruption of *twi* expression in the regions that intersects with the presumptive cephalic furrow, in stage 5 embryos (Fig. 4, B1’- B2’’), when compared to wild-type embryos (Fig. 4, B1-B2). We refer to this region, marked by an arrow, as the DL ‘weak activation region’ and have used the acronym ‘WAR’ to refer to the same. The lack of *twi* activation is not transient and persisted even at later stages of germ band extension (Fig. 4, B3’ vs B3). We observed similar defects in embryos derived from *dl^SCR^/Df* females (Fig. 4, B1’’, B2’’, and B3’’), but their numbers were dramatically and significantly reduced in comparison to *dl/Df* (Fig. 4F). A re-examination of the DL gradient in *dl/Df* and *dl^SCR^/Df* embryos (Fig 4, A1’ and A1’’) confirms that the gradient is normal in that region, pointing to a local failure of DL mediated activation in the WAR.

**Fig. 4.**
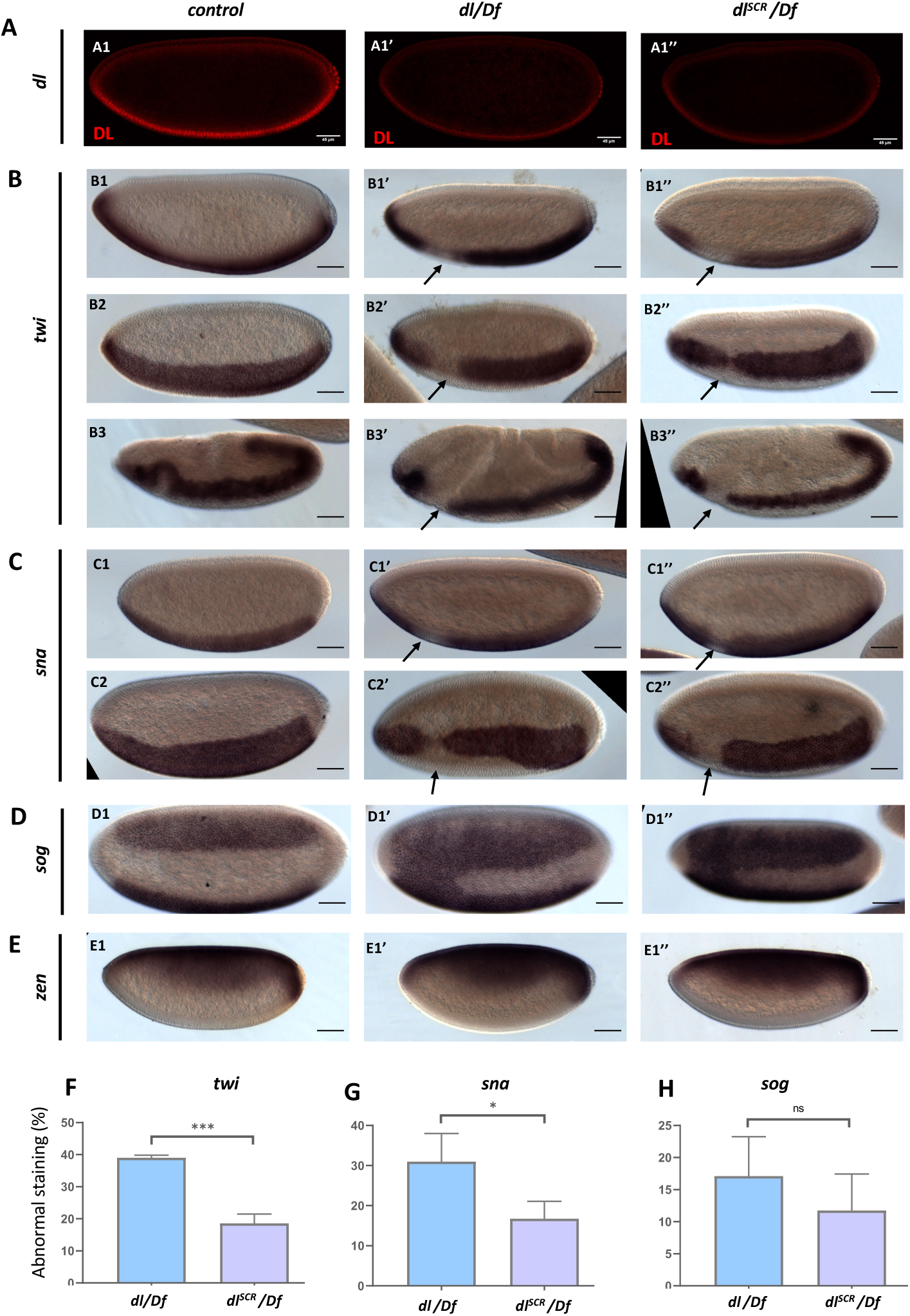
DL activity is altered in the SUMO-deficient mutant. The DL gradient was visualized with a DL antibody in *control*, *dl/Df* and *dl^SCR^/Df* embryos (A). In-situ hybridization images of stage 5 embryos probed with digoxigenin-AP labelled anti-sense RNA probes against *twi* (B), *sna* (C), *sog* (D) and *zen* (E) are shown (B3-B3’’ are stage 7 embryos, an exception). Embryos are oriented with the anterior side to the left and ventral side down (B1-B1’’; B3-B3’’; C1-C1’’; E1-E1’’), or tilted towards the reader (B2-B2’’; C2-C2’’; D1-D1’’), for control (B1-E1), *dl/df* (B1’-E1’) *dl^SCR^/df* (B1’’-E1’’). Arrows indicate a narrowing or an absence of the *twi* (A) and *sna* (B) pattern at the region of the presumptive cephalic furrow. D1’-D1’’ shows a fusion of the *sog* gradient near the ventral cephalic region. Embryos showing a deviation from the normal pattern (narrowing/absence/fusion) for *twi, sna,* and *sog* were plotted as a percentage of total stained embryos, for the wild type, *dl/df* and and *dl^SCR^/df* (F, G, H). ∼50 embryos were scored in each technical replicate, across three replicates. Data represented as mean ± SEM, unpaired t-test, (ns) *P* >0.05, (***) *P* < 0.001, (*) *P* <0.05.

DL and Twi work synergistically to activate *sna* which is critical for mesoderm specification. Haploinsufficiency of *dl* also manifests as a severe loss or absence of *sna* at the WAR, closely mirroring the defects in *twi* expression in *dl/Df* and *dl^SCR^/Df* embryos (Fig. 4, C1- C2’’). While ∼30% of *dl/Df* embryos appear defective in *sna* expression, only ∼15% of *dl^SCR^/Df* embryos display *sna* abnormalities (Fig. 4C1-2, C1’-2’, C1’’- 2’’ and Fig. 4G). In ∼10-15% of *dl/Df* and *dl^SCR^/Df* embryos (Fig. 4, D1’, D’’; Fig. 4G), the *sog* gradient is expanded in the WAR, allowing the two lateral *sog* stripes to fuse ventrally (Fig. 4, D1’ and D1’’). The expansion of *sog* is a direct consequence of the weaker expression of *sna/*Sna in the WAR. *zen*, repressed in the ventral and lateral regions by DL and expressed only at the dorsal-side of the embryo remained unperturbed in the haploinsufficient embryos of both *dl* and *dl^SCR^* (Fig. D1-D1’’). This was in stark contrast to the failure of DL-mediated activation, suggesting that DL-mediated repression was not influenced by the SUMOylation status of DL, even under haploinsufficient conditions.

Thus, the *dl^SCR^* allele rescues the failure of activation in the WAR for a large fraction of haploinsufficient embryos. The data described in this section argues for a role for SUMO conjugation of DL in regulating activation of DL target genes, especially *twi*, in the WAR. SUMO conjugation competent embryos (wild type) have a high failure rate for activation of *twi* and *sna*, in the WAR, under haploinsufficient conditions, when compared to DL^SCR^. The *in-situ* data presented in this section is excellent for discerning spatio-temporal changes in the expression of DL target genes. What is yet unanswered is the effect of DL^SCR^ on the levels of transcripts of DL targets, especially in conditions of haploinsufficiency. For this, we turned to quantitative mRNA measurements using RNA sequencing, described in the next section.

### DL^SCR^ is a stronger transcriptional activator than DL under conditions of haploinsufficiency

In order to obtain a global picture of the transcriptional activity of DL^SCR^, we conducted a quantitative 3’ RNA sequencing experiment on embryos laid by *dl* and *dl^SCR^* mothers, as well as embryos derived from *dl/Df* and *dl^SCR^/Df* mothers. The experiment was conducted at 29 °C for embryos aged 0-2 hours after egg lay, to capture possible quantitative differences in activation of DL target genes on account of maternally deposited DL. Details of the methodology can be found in Materials and Methods. Overall, across the four genotypes (*dl, dl^SCR^, dl/Df* and *dl^SCR^/Df*; Fig. 5A) studied, 194 genes are differentially expressed (−0.58 >log_2_ Fold change> 0.58, at FDR<0.1), visualized as a heat map (Fig. S3A). A gene ontology analysis confirmed significant enrichment of genes encoding proteins involved in DNA-binding and embryonic developmental processes (Fig. S3B). Of these, we focused on genes that are bona-fide DL targets (n=163, Fig. 5B), collated from studies on DV mutants using microarray chips, ChIP-chip analysis and bioinformatics studies (Markstein *et al*. 2002; Stathopoulos *et al*. 2002; Zeitlinger *et al*. 2007). 19 of the differentially expressed genes (DEGs), in our study, are DL targets (Fig 5B), and we generated a heat map to visualize the differences in gene expression across the four genotypes (Fig. 5C).

**Fig. 5.**
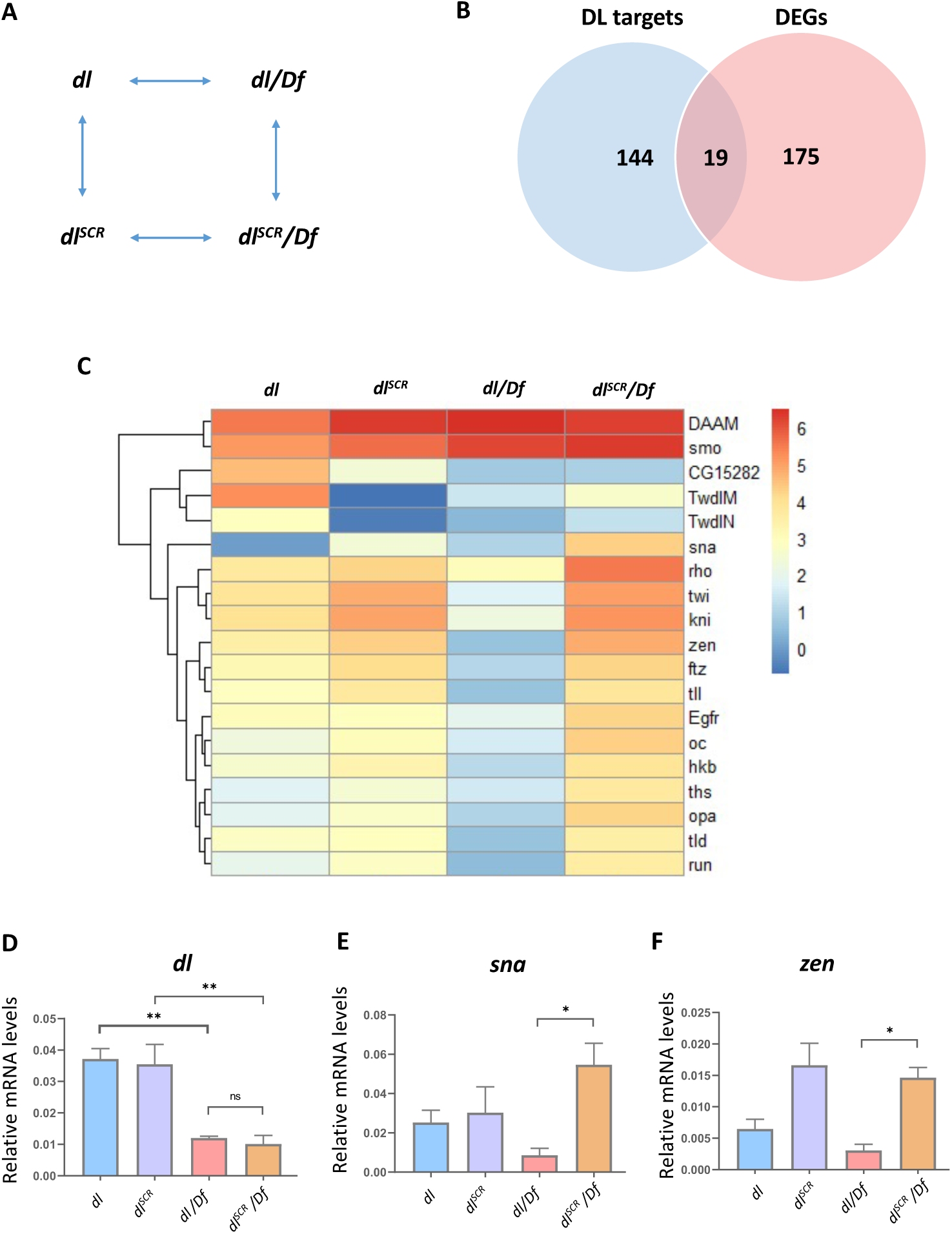
DL^SCR^ displays higher transcriptional activity. Maternal genotypes of the embryos used for the 3’ RNA-seq analysis and their pairwise comparison to obtain differentially expressed genes (DEGs) is presented in (A). Genes that were identified as direct targets of DL from published literature and DEGs across all the conditions are represented as a Venn diagram in (B). The subset of DEGs with known binding sites for DL are represented as a heatmap, for *dl, dl^SCR^, dl/Df,* and *dl^SCR^/Df* embryos at 29 °C, in (C). LogCPM values are plotted. (D), (E) and (F) denote relative mRNA expression levels of *dl, sna,* and *zen* transcripts respectively, measured by qRT-PCR analysis, for 0-2 hour embryos laid by mothers of the indicated genotypes at 29 °C. N=3, mean ± SEM, Ordinary one-way ANOVA, (ns) *P* >0.05, (**) *P* < 0.01, (*) *P* <0.05

As expected, analysis of the transcript levels (represented as log_CPM_ values) clearly indicates that genes like *twi*, *kni*, *zen*, *ftz*, *tll*, and *tld* show lowered transcripts in *dl/Df* compared to *dl*. There is a clear dose response relationship between reduced DL concentrations and reduced transcriptional activation of DL target genes. Exceptions include *Dishevelled Associated Activator of Morphogenesis* (*DAAM*) and *smoothened* (*smo*), whose transcript levels go up significantly, with decreased *dl* levels. When compared to *dl*, *dl^SCR^* did not display statistically significant differences for *twi*, *kni*, *zen*, *ftz*, and *tll*, though average values of transcripts were higher for these and other DL targets (Fig. 5C). Intriguingly, 14 DL target genes are significantly upregulated in *dl^SCR^/Df*, when compared to *dl/Df* (Fig 5C; Fig. S3C). This suggests that for DL^SCR^, the lowered dose of DL in the haploinsufficient embryos is compensated for by higher transcriptional activation of critical DL target genes. These genes include DV patterning targets such as *twi, sna, zen* and *rho* and anterio-posterior (AP) target genes such as *kni* and *hkb* (Fig. S3C). The transcript levels for *dl*, *sna* and *zen* were further independently assessed in a qRT-PCR experiment (Fig. 5 D-F). *dl* transcripts are comparable across *dl/Df* and *dl^SCR^/Df*, downregulated by ∼50% compared to the control (Fig. 5E). *sna* and *zen* however show 6-8 fold higher transcript levels in the *dl^SCR^/ Df*, in agreement with our 3’ mRNA sequencing data (Fig. 5F, 5G). *zen* is a target for DL-mediated repression, but in the absence of expansion of the *zen* expression domain, the transcript data suggests that a transcriptional activator responsible for switching on *zen* may be indirectly influenced by DL.

The greater rate of hatching and survival of *dl^SCR^/Df* when compared to *dl/Df* is possibly a function of increased, compensatory transcription of DL targets, in addition to the previously described rescue of activation in the WAR. SUMOylation of DL thus plays a global role in decreasing transcription of DL targets in general and in addition has a special role in the WAR for activation of *twi/sna*. The higher activation of DL target genes in *dl^SCR^/Df*, leads us to hypothesize that SUMO conjugation of DL may be part of a negative feedback loop to curtail excess transcription of DL targets. SUMOylation of DL is triggered when transcript levels (or protein levels) of one or more DL targets is high and this leads to increase in DL SUMOylation and to attenuation of DL mediated transcription. To understand if the negative regulation of DL targets is a general feature of SUMOylated DL, in the next section, we examine Toll/DL signalling, in host defense, during the larval stages of the animal.

### SUMO restrains DL activity in the larval immune response

Earlier, Bhaskar and co-workers (Bhaskar *et al*. 2002) found that DL^SCR^ is a better transcriptional activator, assessed by luciferase reporter activity on artificial promoter clusters in S2 cells. Since S2 cells are hematopoietic in origin, we reasoned that SUMO conjugation of DL may also influence the immune response in animals. The *dl^SCR^* line allows us to conduct similar experiments in the larvae, with twin advantages over S2 cells of working in the animal and in the absence of confounding wild-type DL in the background. The *Drosophila* larval system has been utilized extensively to understand the physiological roles for Toll/NF-κB signalling in host defense (Sorrentino *et al*. 2004; Matova and Anderson 2006, 2010; Williams 2007; Govind 2008; Schmid *et al*. 2014; Kenmoku *et al*. 2017). Septic injury by gram positive bacteria (or fungal infections) leads to the upregulation of *dl* transcripts and translocation of DL to the nucleus in the larval fat body, a major effector site of the humoral response (Reichhart *et al*. 1993; Lemaitre *et al*. 1995; Manfruelli *et al*. 1999). Similar behavior is exhibited by Dorsal-like immune factor (Dif), a DL paralog (Ip *et al*. 1993; Meng *et al*. 1999; Govind 1999; Hoffmann 2003). Gain of function mutants in Toll/ NF-κB signaling display an over-proliferation of hemocytes (Qiu *et al*. 1998; Matova and Anderson 2006). Studies have also implicated Dif and DL as effectors causing melanotic tumors when constitutively nuclear (Huang *et al*. 2005; Chiu *et al*. 2005).

We reasoned that if DL activity was indeed affected in *dl^SCR^*, it might influence blood cell numbers. In the uninfected state, circulating plasmatocytes are the predominant blood cell type in third instar larvae (Meister 2004). We observed that circulating plasmatocytes remained unchanged in *dl^SCR^* mutants (Fig. 6A). We also looked at crystal cells (Fig 5B1), a platelet-like population of cells important for melanisation and wound healing (Vlisidou and Wood 2015). The readout we used was the activation of the melanization cascade in response to heating/boiling of larvae (Rizki and Rizki 1959; Lanot *et al*. 2001). To our surprise, *dl^SCR^* larvae showed a marked increase in crystal cell numbers (Fig. 6, B2) in comparison to the wild type (Fig. 6, B1). Also striking was the near absence of crystal cells in *dl^1^/dl^1^* animals (Fig. 6, B4), defined in literature as a null allele (Isoda *et al*. 1992). The *dl^1^* allele was sequenced and found to possess an S317N mutation (Fig. S4A) in the rel-homology domain (RHD) (Fig. S4B). DL is phosphorylated at S317, and this modification is essential for its nuclear import in response to Toll signalling in the embryo (Drier *et al*. 1999). We find that this residue is critical for the nuclear import of DL in the fat body as well (Fig. S4, D2’’). A severe reduction in crystal cell number is also evident when another null allele of *dl*, *dl^4^* is used in a trans-allelic combination with *dl^1^* (Fig. 6, B5). The introduction of a single wild type copy of dl with *dl^1^* restores the crystal cell number to levels lower than wild type (Fig. 6, B3). DL has previously been demonstrated to be essential to the phenoloxidase activity cascade (Bettencourt *et al*. 2004). Since our crude readout for crystal cells is its ability to activate the melanization cascade, we are not in a position to state if DL is essential in determining cell fate, or is affecting the phenoloxidase cascade as previously observed. Crystal cell fate is determined by interactions between Serpent, Lozenge and U-shaped (Banerjee *et al*. 2019). Additionally, Rel factors are known to synergize with Srp, activating the immune transcriptional program (Senger *et al*. 2004). Though evidence for DL/Srp co-operativity in determining hematopoietic cell-fate is lacking, we do see a direct correlation between melanized cells and levels of DL activity, with *dl^null^* animals lacking melanization and *dl^SCR^*, our presumptive transcriptionally active allele showing the highest levels.

**Fig. 6.**
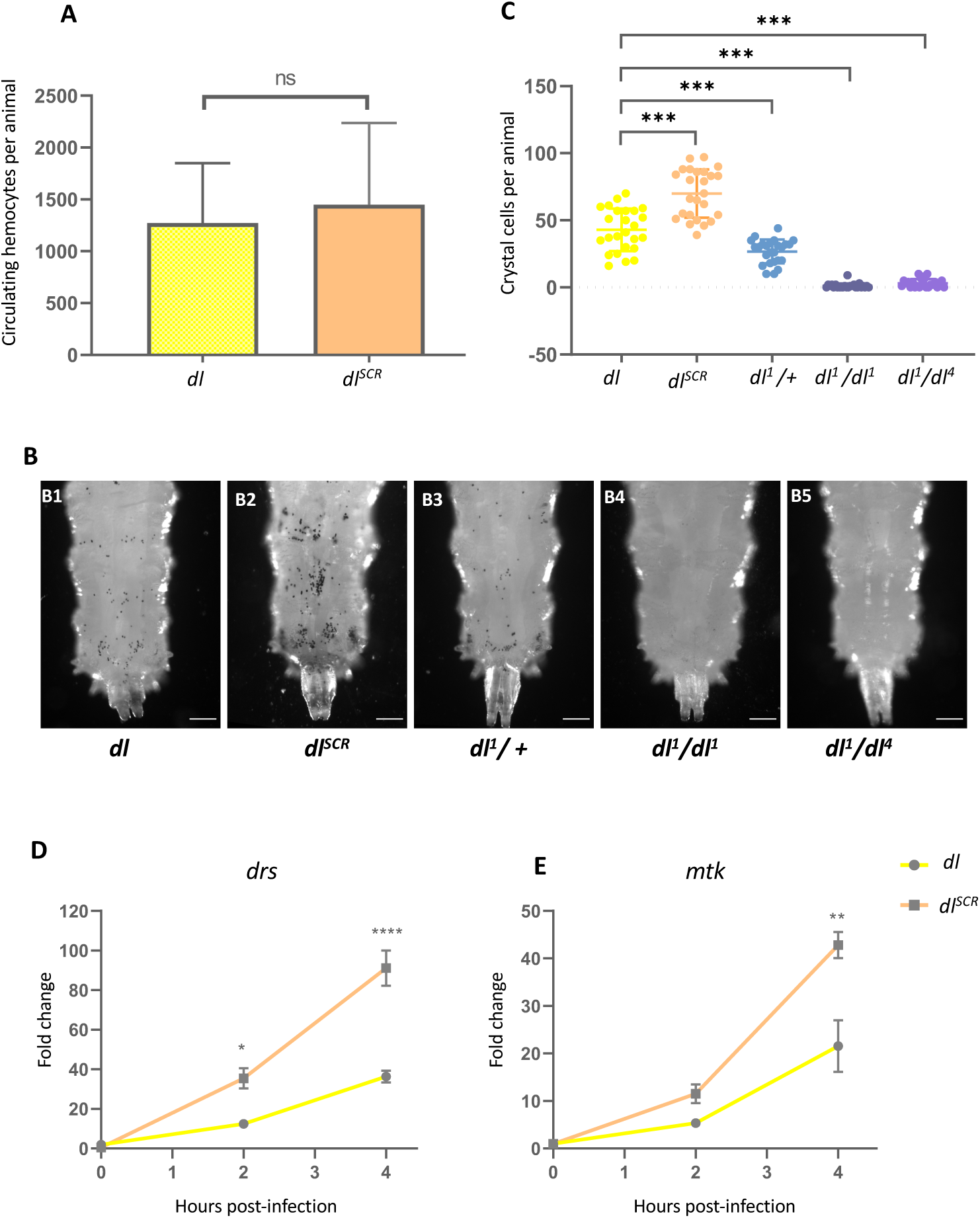
DL^SCR^ is a robust immune effector in the larva. Total circulating hemocytes for *dl* and *dl^SCR^* are plotted as a bar graph in (A). N=3, Mean ± SEM, Unpaired t-test, (ns) *P* >0.05. Crystal cells in the third-instar larva were observed under a bright-field microscope, for the genotypes indicated in (B). The last three posterior segments were imaged with the dorsal side facing the viewer. The number of crystal cells in the posterior segments were counted per animal for each genotype, and are represented in (C). N=3, Mean ± SEM, ordinary one-way ANOVA, (***) *P* < 0.0001. Transcript levels of Toll-responsive AMPs – *drs* and *mtk* (D) analysed by qRT-PCR are plotted for the control and *dl^SCR^*. Data was collected at 0, 2, and 4 hours after septic injury with the gram positive pathogen *S. saprophyticus,* in third instar larvae. N=3, Mean ± SEM, Two-way ANOVA, (*) *P* <0.05, (**) *P* <0.01, (****) *P* <0.0001. Data is representative of at least 8 larvae per replicate, across three independent biological replicates.

We also monitored the temporal expression of AMPs upon septic injury in *dl* and *dl^SCR^*, to gauge the humoral response. Third instar larvae were infected with the gram-positive bacteria *Staphylococcus saprophyticus* and the induction of Toll-specific AMPs *drosomycin* (*drs*) and *metchnikowin* (*mtk*) was analysed using qRT-PCR at 2 hours and 4 hours post-infection. DL^SCR^ induced AMPs to a two-fold higher level, in comparison to DL (Fig. 6D, E). The effect was more prominent at 4 hours post-infection, with both *drs* and *mtk* showing significantly higher expression (Fig. 6D, E). These above results agree with our thesis that DL^SCR^ is a stronger transcriptional activator, and are also in agreement with S2 cell data published previously.

### Dynamics of nuclear import of DL^SCR^ in the larval fat body

In the context of the embryo, DL import appeared to be normal in the SCR allele, with no evidence that supported a change in the DL DV gradient. The primary difference between the DL and DL^SCR^ was seen was in haploinsufficient conditions, where the overall DL-mediated activation was weaker (Fig. 5D) due to low concentrations of DL in the nucleus. In addition to global lowering of transcripts of DL target genes, a complete loss of activation of *twi* was seen in a specific spatiotemporal region, the WAR. In DL^SCR^ embryos, the absence of DL SUMOylation appeared to suppress the weakened transcriptional activation. This is in line with the higher transcriptional activation seen in S2 cells and also in the larval fat body for DL^SCR^. We measured the extent of nuclear import in the fat body of larvae. In un-infected larvae, DL^SCR^ is retained in the cytoplasm, similar to the wild type (Fig 7, A1’’-A2’’). Intensity-based quantitation of the nuclear to cytoplasmic ratio (N/C) is also comparable (Fig. 7B), indicating that SUMOylation of DL is dispensable in terms of retention of the DL/Cact complex in the cytoplasm.

**Fig. 7.**
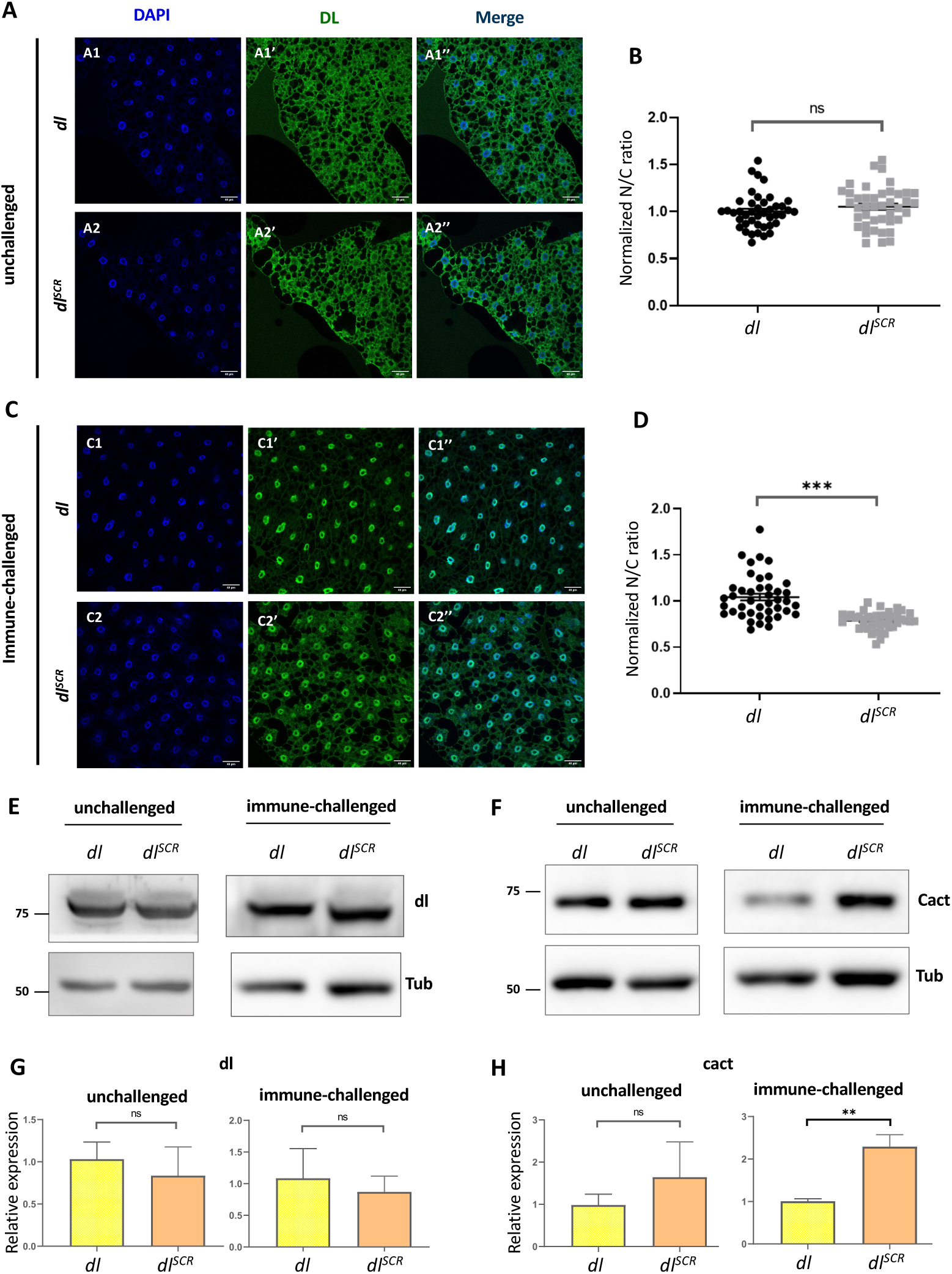
DL^SCR^ is responsive to Toll signalling in the larval fat body. DL is visualized via antibody staining (green), in the uninfected state (A) and infected state (C). Nuclei are labelled with DAPI (blue). Merged images A1’’ and A2’’ indicate uniform distribution in fat body cells. DL levels in the cytoplasm and nucleus were quantified and plotted as a nuclear/cytoplasmic (N/C) ratio for control and mutant (B). The N/C ratio was calculated for >40 cells in at least 5 fat bodies, across three independent replicates. Individual values are represented on a scatter plot, bar denotes mean ± SEM, statistical significance inferred by unpaired t-test, (ns) *P* >0.05, (***) *P* <0.001. DL predominantly partitions to the nucleus 60 minutes after infection with *S. saprophyticus*, evident in merged images C1’’ and C2’’. N/C ratio was quantified and plotted for *dl^SCR^* and *dl* (D), as in (B). Protein levels of DL and Cact in the unchallenged and immune-challenged fat body were determined by a western blot, shown in (E) and (F) respectively. Protein levels were normalized to the loading control (tubulin) and quantified, represented as relative expression levels below the respective blots for *dl* (yellow bar) and *dl^SCR^* (orange bar) (G and H). N=3, bar chart represents mean ± SEM, statistical significance calculated by unpaired t-test, (ns) *P* >0.05, (***) *P* <0.001.

We next monitored the status of DL in the fat body, 60 minutes after an immune challenge with the gram-positive pathogen *Staphylococcus saprophyticus*. DL^SCR^ is competent in its ability to enter the nucleus (Fig. 7, C2’’), at par with the wild-type. Paradoxically, the normalized N/C ratio for DL appears to be lower than wild-type, in the *dl^SCR^* mutants (Fig. 7D), indicating that relative to wild-type, more DL is retained in the cytoplasm or that less DL is imported to the nucleus. The images (Fig. 7C) suggest that the former is true, with a larger fraction of DL^SCR^ apparently retained in the cytoplasm, when compared to DL animals. The retention of DL^SCR^ in the cytoplasm can be because of a number of reasons. There could be (i) an increased affinity of DL^SCR^ for Cact (ii) decreased rate of nuclear import or increased rate of nuclear export or (iii) increase in Cact concentration in the cytoplasm, which would in turn stabilize and retain DL.

Western blots suggest that both DL and DL^SCR^ are expressed at similar levels, both in unchallenged and *Staphylococcus* challenged larvae (Fig. 7E). Intriguingly, Cact, which is degraded on infection induced Toll signalling is more stable in infected conditions, in *dl^SCR^* animals (Fig. 7F). The *cact* genomic region has binding sites for DL, allowing DL to positively regulate Cact levels (Nicolas *et al*. 1998; Paddibhatla *et al*. 2010). Though there are multiple possibilities, we hypothesize that the DL^SCR^ allele, being more active, leads to increased transcription of *cact*. Excess Cact in the cytoplasm, in turn, leads to an efficient retention of DL.

In summary, the increase in stabilized Cact during infection can explain the better cytoplasmic retention of DL^SCR^. DL^SCR^, when available in the nucleus, is more active than wild type DL and leads to a stronger activation of DL targets, such as *cact, drs* and *mtk*.

### A mathematical model to investigate the activity of SUMOylated DL

Data from the embryo, as also the larval cellular and humoral arms all point to heightened transcriptional activation for DL^SCR^. In the embryo, the evidence suggests that SUMO functions to reduce DL-mediated activation globally, with the WAR being a special case where there is an additional local failure of *twi* activation. In the fat body, there is no equivalent to the WAR, but DL target genes are activated more strongly by DL^SCR^, and in addition there is evidence for retention of DL in the cytoplasm, presumably in response to higher levels of Cact in the cytoplasm, which in turn is a result of greater activation of *Cact* by DL. UnSUMOylated DL (DL^U^) appears to be a stronger transcriptional activator and SUMOylation of DL a mechanism to attenuate DL mediated activation. Since levels of SUMOylated DL (DL^S^) are estimated to be <5% in the best case, it is difficult to examine this species experimentally. One way to gain additional insight into roles for the DL^S^ species, would be to use a mathematical model and evaluate numerically the effect of DL SUMOylation.

Here, we model the effect of DL^S^, which a small fraction of DL in the cell, with DL^U^ being the predominant species. For modelling, we used previously reported association and transport parameters (Tay *et al*. 2010) as an approximation. We assumed that the total DL and the fraction that is SUMOylated remains unchanged in the timescale considered for the simulation. The effect of SUMOylation might be to change the extent of binding to Cact, partitioning between the nucleus and cytoplasm, promoter-binding affinity, or ability to enhance transcription (Fig. 8A). We have derived equations, as listed in Materials and Methods, whose numerical solution enables us to calculate the intensity of gene expression in DL^WT^ and DL^SCR^ as a function of the various affinities, partition coefficients, and total amounts of Cact, DL^S^, DL^U^ and binding sites. Using previously reported parameters, we simulated the system for various hypothesized effects of SUMOylation on thermodynamic parameters.

**Fig. 8.**
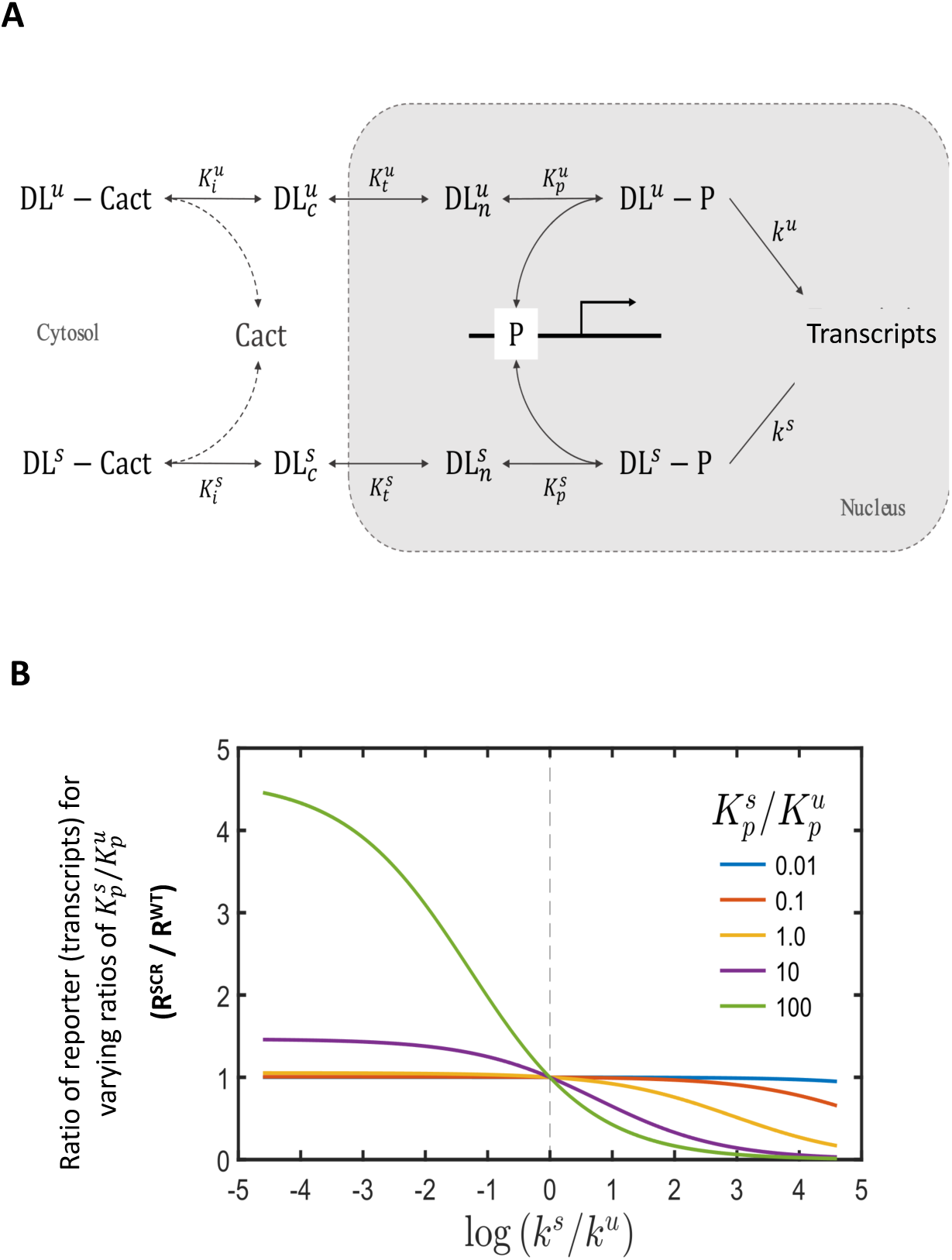
Mathematical model to understand roles for SUMOylated DL. SUMO conjugated DL in the cell is <5% of total DL, even under conditions of stress such as high temperatures. We utilize a mathematical model (A) to understand conditions/parameters where a small fraction of SUMOylated DL (DL^S^), fixed at 5%, can influence Toll signalling. The cytosolic and nuclear species are denoted as DLC and DLN respectively, while unSUMOylated DL is DL^U^. P represents the binding site for DL^S^ and DL in the nucleus. Ki, Kt and Kp denote the equilibrium constants for DL:Cact association, transport of DL into the nucleus, and binding of nuclear DL to the promoter P, respectively. (B). Simulating the effect of SUMOylation of DL on transcription of DL target genes. The reporter expression levels for the DL^SCR^ is greater than the corresponding WT level when the specific transcription rate is lower (*k*^s^/*k*^u^ <1) for the SUMOylated DL relative to unSUMOylated DL. Tighter binding to promoter 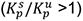 enhances this effect.

Model simulations (Fig. 8B, Fig. S5) suggest that change in extent of activation, and not in binding affinity or N/C partitioning, leads to significant differences in the reporter transcript in DL^WT^ vs DL^SCR^. Further, the simulations suggest that the observed effect of SUMOylation on transcription of target genes is primarily due to the weaker ability of DL^S^ to promote transcription. This effect is modulated by other factors such as inhibition by Cact, partitioning into nucleus and binding to DL binding sites on the promoter. For instance, stronger binding of DL^S^ to the promoter site coupled with weaker transcriptional activity of the bound DL^S^ results in higher reporter expression in DL^SCR^ relative to DL^WT^. Hence in the case of DL^WT^, DL^S^ is bound to the promoter site, resulting in lower transcription due to its weaker (or null) activity compared to DL^U^. When SUMOylation is abrogated, the DL^U^ binds and its higher specific transcriptional activation results in increased expression. This may also provide an explanation for the ability of DL^SCR^ in the embryo to rescue the effect of *dl* haploinsufficiency. Lower total DL due to loss of an allele may lead to lower activation, which increases to near WT levels when SUMOylation is abrogated.

## Discussion

Comprehensive proteomic studies across species have led to the identification of SUMO conjugated proteins (Wykoff and O’Shea 2005; Handu *et al*. 2015; Hendriks and Vertegaal 2016; Pirone *et al*. 2017; Hendriks *et al*. 2018). Overall, roles for SUMO conjugation of a substrate protein appears to be context dependent, with its effect on the target not easily predicted. One class of proteins studied is transcription factors (TFs) (Verger *et al*. 2003). TFs are major downstream effectors of cellular signalling, and can undergo extensive post-translational modifications. Nearly 50% of DNA-binding TFs are SUMOylated in humans (Hendriks *et al*. 2017). Upon SUMOylation, changes in the transcriptional output of TFs can be brought about by alterations in DNA-binding, eviction from the chromatin, or a re-shaping of their protein-interaction landscape (Ouyang and Gill 2009; Raman *et al*. 2013; Wotton *et al*. 2017; Rosonina *et al*. 2017; Rosonina 2019). In terms of absolute numbers, SUMO is known to predominantly repress TF activity (Müller *et al*. 2000; Poukka *et al*. 2000; Bies *et al*. 2002; Eloranta and Hurst 2002; Abdel-Hafiz *et al*. 2002; Ross *et al*. 2002; Sapetschnig *et al*. 2002; Nishida and Yasuda 2002; Tian *et al*. 2002). Amongst the well-studied TFs is the NF-κB family (Kracklauer and Schmidt 2003; Mabb and Miyamoto 2007). SUMOylation of NF-κB was first demonstrated in *Drosophila* by the Courey lab, with DL being a target of SUMOylation (Bhaskar *et al*. 2000). Recently, Relish SUMOylation has been shown to be important to restrain its auto-activation (Tang *et al*. 2021). In mammals, RelA (p65) undergoes SUMOylation after TNFα stimulation, aided by the E3 ligase PIAS3. In addition to identifying a role for SUMOylation in RelA repression, the authors also observe, interestingly, that only the DNA-bound form of RelA is SUMO-modified (Liu *et al*. 2012). RelB, another NF-κB family transcription factor, is also negatively regulated by SUMOylation, though its DNA binding remains unchanged (Leidner *et al*. 2014). The authors speculate that a repression of Rel activity by SUMOylation can be attributed to the assembly of transcriptional repressors at the DNA, via their SIMs.

In *Drosophila*, the Toll/NF-κB cascade has been best studied in two diverse contexts, in DV patterning (Fig. 9A) and in the immune response (Fig. 9B). In early development, where perturbations to the nucleo-cytoplasmic gradient of DL could derail DV patterning, we do not see any effect of lack of SUMOylation on DL activity across the DV axis. Increase in the transcriptional activity of DL^SCR^ becomes apparent only in conditions of haploinsufficiency (*dl^SCR^/Df*), where DL target genes are, surprisingly, activated at wild-type levels. This enhanced transcriptional activation with reduced DL dosage suggests that SUMO conjugation is linked to a negative-feedback loop (Fig. 9C). We visualize the feedback loop as a circuit (Fig. 9C), where excess transcription is sensed by a hypothetical sensor, that leads to SUMOylation of DL by Ubc9. SUMOylated DL, when bound to the promoter, blocks activation by DL. The sensor would ideally sense high transcript levels of DL target genes. miRNAs, known to dampen Toll signalling in the immune-context, may also serve as sensors of DL-activated targets (Li *et al*. 2021). Under conditions of haploinsufficency (*dl/Df*), the sensor is not in play, as it responds only to high levels of DL activity. Hence we see a dose response relationship; a low amount of DL protein leads to lower levels of DL-target transcripts. In the case of *dl^SCR^*, the circuit is broken, with Ubc9 unable to SUMOylate DL^SCR^. DL^SCR^ does not show significantly enhanced activation of DL target genes, as compared to DL, suggesting that maximum transcriptional rates have been achieved with two copies of DL. In the case of dl^SCR^/Df, the circuit is also broken, and as a result, we see higher transcription rates, in the absence of SUMOylated DL. A special feature of haploinsufficiency in the embryo is the WAR. This phenomenon goes beyond a generic lowering of *twi* activation, and is possibly related to the modulation of a physical interactor of DL (Fig. S6) as a function of SUMO conjugation of DL. Previous literature on DL suggests a few possible candidates. These include Daughterless (Da), Achaete-scute complex (AS-C), Nejire(Nej)/CREB binding protein(CBP) and TBP-associated factors (TAFs). Embryos laid by *nej^3^/+*;*dl^1^/+* mothers showed weaker *twi* expression and a lack of expression in the WAR (Akimaru *et al*. 1997). Similar effects on *twi* expression was seen in eggs laid by *dl^1^/+*;*TAF_II_110/+* or *dl^1^/+*;*TAF_II_60/+* mothers (Zhou *et al*. 1998) as also *dl^8^/+*;*da^11B31^/+* mothers (González-Crespo and Levine 1993). We hypothesize that DL forms complexes with these proteins to activate *twi* and this complex formation is especially important in the WAR (Fig. 9D). DL^U^ is efficient at interacting with one (or all) of these proteins whereas DL^S^ de-stabilizes the activation complex. Thus in the *dl^SCR^* animal, the absence of DL^S^ leads to a robust activation of *twi* in the WAR, suppressing the lethality due to haplo-insufficiency. DL functions as a dimer to activate transcription. The low levels of DL^S^ suggest that the existence of homo-dimers of DL^S^ would be a very small fraction of total DL. How then, do hetero-dimers of SUMOylated and unSUMOylated DL bring about a potent impediment of transcription? The answer could lie in a putative conformational change brought about by SUMOylation disallowing the recruitment of cognate co-activators, rendering the dimer non-functional for transcriptional activity, while leaving dimerization properties unaffected.

**Fig. 9.**
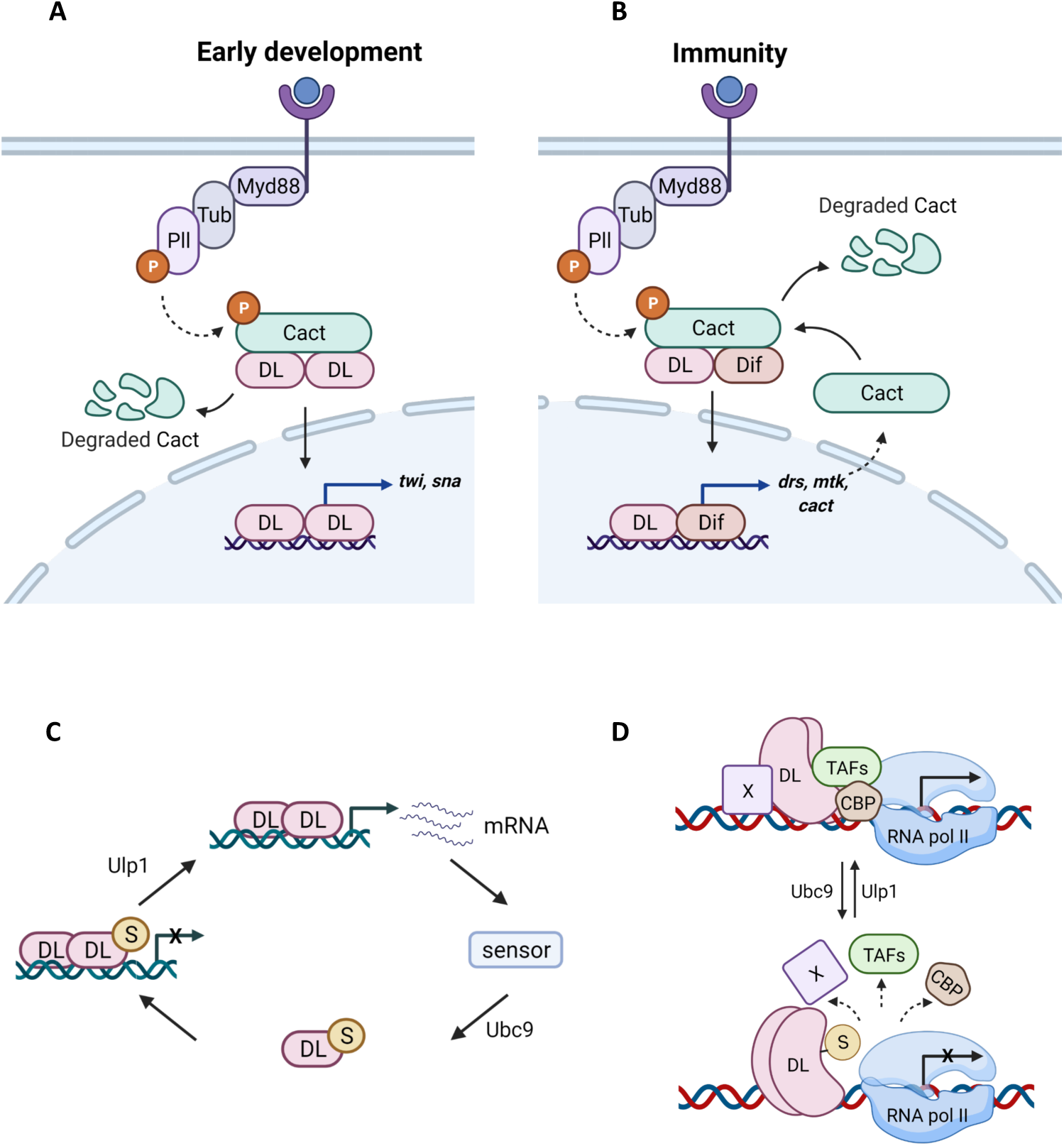
DL SUMOylation attenuates Toll signalling. A probable model for dampening of the Toll signal upon DL SUMOylation. When Toll signalling is initiated, DL migrates to the nucleus, and activates target genes, in both the developmental and immune context (A and B respectively). Once optimum levels of target transcripts are reached, or under conditions of stress, SUMOylation of DL is triggered, through as yet unknown mechanisms, curtailing excessive transcription (C). Transcriptional activity of DL may be regulated via conserved interactions with CBP and/or TAFs in association with with a protein X, most likely GATA-factor Srp in immunity or bHLH proteins like daughterless/achate-scute, in early development (D). SUMOylation of DL may perturb these interactions, attenuating transcription.

Does our data suggest any role for DL in DV patterning under ambient conditions with normal DL concentrations? We hypothesize that DL SUMOylation could fine-tune transcription rates of DL target proteins in response to environmental fluctuations, a normal feature for cold-blooded animals. DL may influence the robustness of the DL gradient, supporting the progression of embryogenesis, for example, in varied range of temperatures. This leads us to suggest that SUMO conjugation of DL is a mechanism for developmental canalization, as hypothesized by Conrad Waddington (Waddington 1959). In the context of the developing fly embryo, a signature for canalization would be mechanisms that would lead to a definite end result, which is embryonic hatching and normal DV patterning, by suppressing the variations that would lead to the reaction deviating from its normal path. SUMO conjugation of DL is thus an excellent example of a post-translational modification that leads to canalization.

In response to infection (Fig. 9B), *dl^SCR^* larvae exhibit an increase in the transcription of Toll-specific AMPs and show higher number of crystal cells. Similar to data seen in the developmental context, SUMOylation of DL attenuates transcriptional activation of DL target genes. *dl^SCR^* animals show 2-4 higher transcripts of *drs* and *mtk*, under infective conditions and 2-fold increase in crystal cells, in the absence of infection. Again, as suggested earlier (Fig. 9C), SUMOylation of DL may be a mechanism to attenuate DL-mediated activation. In addition to co-activators such as CBP or Dlip3 (Ratnaparkhi *et al*. 2008), the GATA-family transcription factor Serpent (Srp) may be an important player. DL, Dif, and Relish are known to synergize with Srp (Petersen *et al*. 1999; Senger *et al*. 2004) in the larvae and SUMOylation of DL may weaken or break these interactions. Srp, along with the RUNX-factor Lozenge (Lz) is critical for specifying crystal cell fate, in embryonic and larval stages (Fossett *et al*. 2003). Our observation that DL^null^ animals have few or no melanised crystal cells while DL^SCR^ larvae have increased crystal cells may point to a hitherto unknown function of DL carried out in assistance with Srp/Lz. The fact that Dif is unable to compensate for DL function in this aspect is also intriguing, highlighting non-redundant, divergent functions for DL and Dif in the larva. Further, In the larval fat body, nuclear partitioning of DL is affected in *dl^SCR^* animals, in response to septic injury. Cact is more stable in the cytoplasm of *dl^SCR^* animals, and this would lead to retention of DL. Nevertheless, transcript levels of DL target genes are higher, leading us to hypothesize that *dl^SCR^* animals have higher *Cact* levels, and higher Cact levels could explain the enhanced retention of DL in the cytoplasm, though the possibility of increased binding affinity of DL^U^ for Cact cannot be ruled out.

Our work further highlights the intricate fine-tuning that regulates signalling cascades. Toll signalling is modulated at multiple levels. Extracellular feedback exerted by serine hydrolase cascades serves as an initial checkpoint for receptor activation. Cact and WntD (Ganguly *et al*. 2005; Gordon *et al*. 2005) act as intracellular, cytoplasmic gatekeepers of DL activation. An additional phosphorylation step is necessary for the nuclear import of DL. Once in the nucleus, DL can interact with partner activators and co-repressors to calibrate the transcriptional output. Our data suggests an additional layer of control, within the nucleus, with SUMO conjugation as a means of keeping DL in check, downstream of its nuclear import. The SUMO conjugation machinery resides in the nucleus and this is the most probable site for SUMO-conjugation/deconjugation of DL. The SUMO conjugase Ubc9 is a physical interactor of DL (Fig. S6) and presumed to be placed proximal to the site of transcription (Bhaskar *et al*. 2000). The SUMO deconjugase Ulp1 may also be similarly localized (Anjum *et al*. 2013). Since a very small proportion of total DL is SUMO conjugated, SUMOylation/deSUMOylation may be a dynamic process that defines the occupancy of ‘active’ DL for transcription. Since DL^U^ is a better transcriptional activator, SUMOylation of DL may be a general mechanism to reduce occupancy of DL^U^ at the promoter regions. Our mathematical model suggests that DL^S^, which is <5% of total cellular DL, binds to the promoter and blocks access to DL^U^, thus attenuating transcription. Additionally, DL^S^ may be deficient in its ability to interact with the core transcriptional machinery or with partner basic helix-loop-helix proteins. DL^S^ is, in all probability, a non-functional variant of DL. Alternatively, though not directly supported by our data, is the possibility that DL^S^, when bound to DNA can recruit a repressor and subsequently lead to deacetylated chromatin that is resistant to transcription.

In summary, in this study we have generated a genome edited *dl^SCR^* line. Using this reagent, we have evaluated roles for SUMO conjugation of DL and found that SUMOylation acts to reduce transcriptional activation of DL target genes in both developmental and immune contexts. There are possibly context specific differences in mechanisms of transcriptional attenuation in the scenarios that we have studied, and these are dependent on proteins that interact differentially with DL^U^ and DL^S^. SUMO mediated attenuation of DL activity thus adds another layer to the complex regulation of Toll/NF-κB signalling.

## Materials and Methods

### Fly husbandry and stocks

Flies were raised on standard cornmeal agar at 25 °C unless stated otherwise. For the *dl* deficiency experiments, flies were crossed and maintained at 29 °C. The following fly stocks were procured from the Bloomington *Drosophila* Stock Centre: *dl^1^/CyO* (3236), *dl^4^/CyO* (7096), and *vasa-Cas9* (51323). The *dl* deficiency allele *w-, y; J4/CyO* containing a precise deletion of the *dl* and *dif* loci, was a kind gift from the Govind laboratory, CUNY, NY.

### Generation of transgenic CRISPR lines

The Fly CRISPR Optimal Target Finder was used to design the gRNA with zero predicted off-target effects. The gRNA sequence 5’ GAAACATACCGCCCATTAAAA 3’ was incorporated into the forward primer sequence GAAACATACCGCCCATTAAAAGTTTTAGAGCTAGAAATAGC. A reverse primer of the following sequence was used: GAAGTATTGAGGAAAACATA. The gRNA was cloned into the pBFv-U6.2 vector as described previously (Kondo and Ueda 2013), using the primers listed above. The 100-mer ssODN sequence is as follows: TTTAACTAGGTTTTTTTTTTGTAGTTTTAGTGTATAAAACTCACCTCTTGGTTCCGTTCGAATGGGCGGTATGTTTTGTGTATTCCAGCAATTCATGTTA

A total of 620 *vasa-Cas9* embryos were co-injected with the gRNA and ssODN, at the C-CAMP facility, NCBS. 450 F0 adults that emerged were crossed with *Tft/CyO* balancer flies individually. Three emergent flies from each cross were balanced further, with the *Tft/CyO* balancer, and maintained as separate lines. Homozygous flies from these founder lines were screened for the presence of the mutation by PCR followed by restriction digestion. For the isolation of genomic DNA, flies were placed in 0.2mL tubes individually and lysed in 50μL of squishing buffer (10mM Tris-Cl pH 8, 1mM EDTA, 25mM NaCl, and 0.2 mg/mL Proteinase K). After incubation at 37°C for 30 minutes, Proteinase K was inactivated by heating at 85°C. 1μL of the genomic DNA was used in a 10 μL PCR reaction. The following primers were used for the PCR: F: CAGTTCTGAGTAAGTCTTTATCGGAGTTCA; R: CCAAAGGGTTGTGGCGAGGTAT. The PCR product was digested with the restriction enzyme BstBI and resolved on a 1.2% agarose gel. Four transformants were obtained after screening 200 lines.

### Cuticle preparation

Embryos were collected for three hours and aged for 22 hours at 25°C or 29°C, depending on the nature of the experiment. They were dechorionated in a 4% sodium hypochlorite solution for 2 minutes. Dechorionated embryos were washed thoroughly under running tap water and transferred to a scintillation vial containing 1:1 methanol: heptane. The vial was shaken vigorously for a few minutes, and de-vitellinized embryos in the lower methanol phase were transferred to a new vial with fresh methanol. Embryos were transferred onto a slide, mounted in 85% lactic acid, and incubated overnight at 55°C on a slide warmer. Cuticles were imaged on a Zeiss Axio imager Z1 microscope, using dark field illumination, with a 10X objective.

### Antibody staining

0-3 hour embryos were dechorionated in 4% sodium hypochlorite for 2 minutes. Embryos were rinsed and fixed in a 1:1 solution of 4% formaldehyde in 1X Phosphate-buffered saline (PBS): heptane for 20 minutes. The aqueous phase containing formaldehyde was removed, and embryos were devitellinized by adding an equal volume of ice-cold methanol followed by vigorous shaking. Devitellinized embryos were washed thrice in methanol. Embryos were re-hydrated and permeabilized by giving six 15-minute washes in 1X PBS containing 0.3% Triton X-100 (0.3% PBS-T). After blocking with 2% Bovine serum albumin (BSA) in 0.3% PBS-T, embryos were incubated overnight at 4°C with the primary antibody. Following four 15-minute washes with 0.3% PBS-T, embryos were incubated with the secondary antibody for an hour at room temperature. Embryos were washed thrice in 0.3% PBS-T, and DAPI was added in the penultimate wash. Embryos were mounted in SlowFade Gold mountant (Invitrogen) and imaged on a Leica Sp8 confocal microscope under a 20X oil-immersion objective. The antibodies used were: Mouse anti-Dorsal, 1:1000 (DSHB 7A4-c) and goat anti-mouse Alexa568 secondary antibody, 1:1000 (Invitrogen).

### RNA in-situ hybridization

Embryos were collected and aged at 29°C. Anti-sense digoxigenin-labelled RNA probes for *twi*, *sna*, *sog,* and *zen* were used and hybridization was carried out as previously described (Tautz and Pfeifle 1989). Anti-Digoxigenin-Alkaline phosphatase antibody (Merck) was used at a concentration of 1:2000 and NBT/BCIP (Merck) was used as the colour-development substrate for AP. Images were acquired on a Zeiss Axio imager Z1 microscope, using DIC optics, with a 10X objective.

### Western blots and their analysis

Fat bodies (8-10 per sample) were dissected in ice-cold PBS and crushed in lysis buffer (2% SDS, 60 mM Tris-Cl pH 6.8, and 1X PIC). Samples were cleared by centrifuging at 21,000*g* for 30 minutes. Total protein was estimated by BCA assay (Pierce) and samples were boiled in 1X Laemmli buffer. Equal amounts of protein (30-40 μg/sample) were resolved on a 10% polyacrylamide gel and transferred onto a PVDF membrane (Immobilon-E, Merck). The membrane was blocked with 5% milk in TBS containing 0.1% Tween20 (TBS-T) for an hour followed by incubation with the primary antibody diluted in 5% milk in TBS-T. Following three washes with TBS-T, the membrane was incubated with the secondary antibody diluted in 5% milk in TBS-T for an hour, at room temperature. The membrane was washed thrice with 0.1% TBS-T, incubated with Immobilon Western Chemiluminescent HRP substrate (Merck), and visualized on a LAS4000 Fuji imaging system. The following antibodies were used: Rabbit anti-Dorsal, 1:5000 (kind gift from the Courey laboratory); Mouse anti-Cactus, 1:100 (DSHB 3H12); Mouse anti-α-Tubulin, 1:10000 (T6074, Sigma-Aldrich); Goat anti-rabbit HRP and Goat anti-mouse HRP secondary antibodies, each at 1:10000 (Jackson ImmunoResearch).

### Microbial infection

*Staphylococcus saprophyticus* (ATCC 15305) was used for the septic injury experiments. For larval infection, the bacteria were grown overnight, concentrated by centrifugation, and the pellet washed with PBS. Larvae were placed on a cold agar plate and infected at the posterior region with a fine insect pin dipped in the concentrated culture, as described previously (Kenmoku *et al*. 2017). Infected larvae were transferred to a fresh sugar-agar plate, at 25 °C and processed at the appropriate time points.

### Fat body staining

Fat bodies from wandering third instar larvae were dissected in ice-cold PBS and fixed in 4% formaldehyde in PBS, for 20 minutes. The tissue was permeabilized by washing thrice in 0.1% PBS-T followed by blocking in 2% Bovine serum albumin (BSA) in 0.1% PBS-T. The tissue was incubated overnight with the primary antibody diluted in 2% BSA in 0.1% PBS-T, at 4°C. Following three 15-minute washes with 0.1% PBS-T, secondary antibody diluted in 2% BSA in 0.1% PBS-T was added and incubated for an hour at RT. After three 15-minute washes with 0.1% PBS-T, with DAPI being added in the second wash, the tissue was mounted in SlowFade Gold mountant (Invitrogen) and imaged on a Leica Sp8 confocal microscope under a 20X oil-immersion objective. The antibodies used were: Mouse anti-Dorsal, 1:1000 (DSHB 7A4-c) and goat anti-mouse Alexa488 secondary antibody, 1:1000 (Invitrogen). Mean pixel intensity for DL staining in the cytoplasm and the nucleus was quantified using ImageJ software. The cytoplasmic intensity was averaged across three circular ROIs per cell and the same ROI was used to calculate the nuclear intensity. 5-7 cells per fat body were analysed for at least 7-9 fat bodies across three biological replicates.

### Quantitative PCR

RNA was extracted from appropriately staged embryos or whole larvae (n=10 /sample) using the RNeasy Plus Universal mini kit (Qiagen) according to the manufacturer’s instructions. 1 μg of total RNA was used to generate cDNA using the High-Capacity cDNA Reverse Transcription kit (Thermo Fisher Scientific). The qPCR reaction was performed on a qTOWER^3^ real-time thermal cycler (Analytik Jena) with KAPA SYBR FAST master mix (Sigma-Aldrich). Gene expression was monitored using gene-specific primers. Transcript levels were calculated using the comparative Ct method to obtain fold change values. Relative mRNA levels were calculated using the delta Ct values. Rp49 was used as a reference gene. The following primer pairs were used (Forward primer, F and reverse primer, R):

*rp49* F: GACGCTTCAAGGGACAGTATC, *rp49* R: AAACGCGGTTCTGCATGAG;

*dl* F*: ATCCGTGTGGATCCGTTTAA, dl* R: AATCGCACCGAATTCAGATC;

*twi* F: AAGTCCCTGCAGCAGATCAT, *twi* R: CGGCACAGGAAGTCAATGTA;

*sna* F: CGGAACCGAAACGTGACTAT, *sna* R: CCTTTCCGGTGTTTTTGAAA;

*zen* F: TACTATCCAGTTCACCAGGCTAA, *zen* R: TCTGATTGTAGTTGGGAGGCA;

*mtk* F: GCTACATCAGTGCTGGCAGA, *mtk* R: TTAGGATTGAAGGGCGACGG;

*drs* F: CTGTCCGGAAGATACAAGGG, *drs* R: TCGCACCAGCACTTCAGACT

### Quantitative RNA sequencing and analysis

Cages containing flies of the appropriate genotype were set up with sugar-agar plates. Plates were changed twice after 1 hour intervals and the third collection was used for the experiment. Embryos were collected at 29 °C for two hours. RNA was isolated from two biological replicates for each sample using the RNeasy Plus Universal mini kit (Qiagen). RNA concentration was determined using a NanoDrop instrument, and 500ng of RNA was used to generate the cDNA library with the QuantSeq 3′mRNA-Seq Library Prep Kit FWD for Illumina (Lexogen), according to the manufacturer’s instructions. The library size and quality were determined on a Bioanalyzer with a high sensitivity chip (Agilent) and concentration assessed using a Qubit fluorometer, with a dsDNA High Sensitivity assay kit (Thermo Fisher Scientific). The equimolar, pooled library was sequenced on an Illumina NextSeq 550 system, generating 75 bp single-end reads. The sequencing files obtained were uploaded onto BlueBee’s genomics analysis platform (https://lexogen.bluebee.com/quantseq/). Reads were trimmed in BlueBee using bbduk (v35.92). Reads were aligned, counted, and mapped using BlueBee’s STAR-aligner (v2.5.2a), HTSeq-count (v0.6.0), and RSEQC (v2.6.4), respectively. A DESeq2 application within BlueBee (Lexogen Quantseq DE 1.2) was used to obtain normalized gene counts and identify differentially expressed genes based on a false discovery rate (FDR) cut-off *P-*adjusted value <0.1. Downstream analysis was performed on EdgeR. Raw count data was transformed using the logCPM function to obtain values for the heatmap, generated using pheatmap in RStudio. GO enrichment analysis was performed using gProfiler (https://biit.cs.ut.ee/gprofiler/gost).

### Blood cell preparation and counting

Third instar larvae were cleaned with copious amounts of water and a brush, and placed individually in a drop of 20 μL ice-cold PBS (5 per replicate) on a clean glass slide. Larvae were carefully ripped open in PBS using watchmaker’s forceps, without damaging the internal organs. The carcass was discarded, and 10 μL of the PBS solution containing blood cells was transferred to a Neubauer hemocytometer chamber (Hausser Scientific). Plasmatocytes were counted on a Zeiss Axio Vert.A1 microscope at 40X magnification using phase-contrast optics. To visualize crystal cells, wandering third instar larvae (8 per replicate) were heated at 60°C in a water bath for 10 minutes. Images were acquired on a Zeiss Axio Vert.A1 microscope at 10X magnification. Crystal cells in three terminal segments were counted and plotted.

### Statistical Analysis

All experiments were performed in three biological replicates, unless stated otherwise. Data is presented as mean ± SEM. Statistical analysis was performed using GraphPad Prism8.

### Mathematical modelling and simulation

The effect of DL on reporter gene expression was modelled as depicted in Suppl. Fig. 5. DL binding to Cact, transport to the nucleus, and binding to the promoter site was modelled as reversible reactions with kinetics given by the law of mass action. Rate (R) of reporter gene expression was modelled as a first order process dependent on the bound promoter level. The reaction rate constants for all individual processes involving DL^S^ Relative rates can also be interpreted as relative steady state reporter amounts if it assumed that the reporter degradation is first order. DL^S^ is assumed to participate in all reactions but with different specific rate constants relative to DL^U^. It is assumed that in the time interval being considered, total protein amounts are conserved and free and bound protein concentrations are at a (pseudo) steady state. A mass balance on individual proteins and complexes results in the following set of equations (Eq. 1-9).

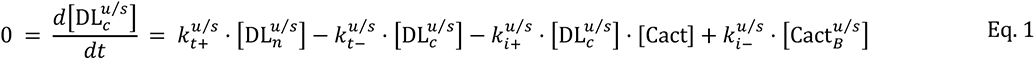

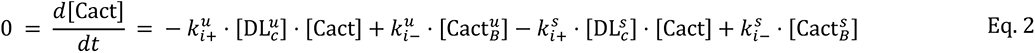

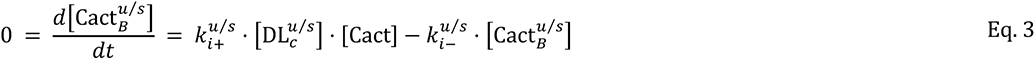

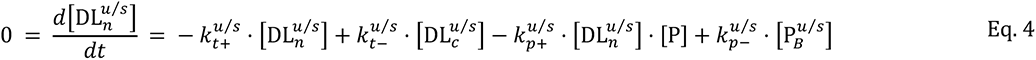

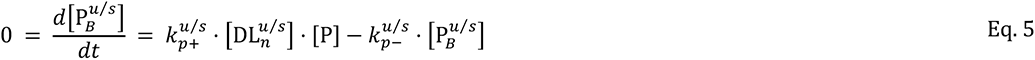

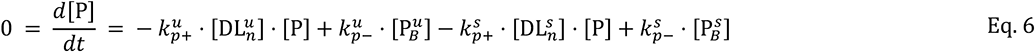

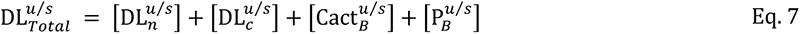

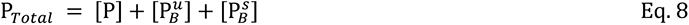

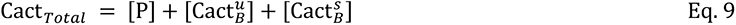

The algebraic expressions in combination with conservation of total DL, binding sites and Cact are used to solve for 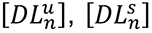 as functions of total DL concentrations. This gives the following two equations (Eq. 10-11) for the wild type,

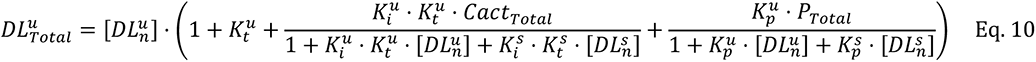

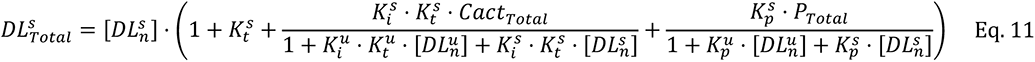

where 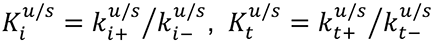, and 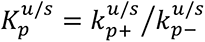. These equations are numerically solved using reported parameter values (Fig. S5; (Tay *et al*. 2010)) and the rate of reporter expression is calculated using

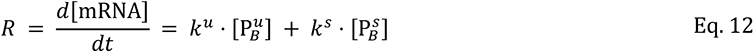

For WT, the total DL reported by Tay et al. is assumed to comprise of 95% unSUMOylated and 5% SUMOylated DL; whereas for DL^SCR^, the entire DL is assumed to be unSUMOylated 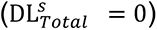. That is,

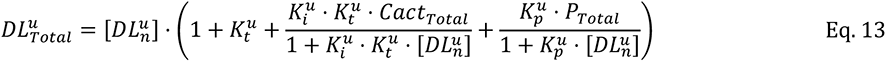

Similar to WT, the above equation is also numerically solved using the *vpasolve* function in MATLAB, with the rate of reporter gene expression having contributions only from unSUMOylated DL 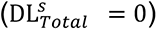. We assume that the effect of SUMOylation is to change the values of the specific reaction rate constants for reactions involving DL. For instance the specific transcription rate (*k^S^*) when promoter is bound to DL^S^ might be higher or lower than the rate (*k^u^*) when promoter is bound to DL^U^. The relative reporter expression is calculated at different values of the relative reaction rate constants for DL^S^ and DL^U^ by solving equations with rate constants set at each of those values.

## Contributions

SH and GR conceptualized the project and designed the experiments. SH performed all the experiments. AS and CG generated the mathematical model and AS carried out the numerical calculations. SH, AS, CG and GR analysed the data and wrote the manuscript. GR supervised the project and acquired funding.

## Acknowledgements

We thank: Bloomington *Drosophila* Stock Center (BDSC), Indiana, supported by NIH grant P40OD018537, for fly stocks; Fly facility at the National Centre for Biological Sciences (NCBS), Bangalore for embryonic injections. Dr. Deepti Trivedi for her input on design of the CRISPR experiments; IISER Pune microscopy facility; Snehal Patil and Yashwant Pawar for fly media and stock maintenance. LS Shashidhara, Girish Deshpande and Richa Rikhy for helpful discussions.

## References

Abdel-Hafiz H., G. S. Takimoto, L. Tung, and K. B. Horwitz, 2002 The inhibitory function in human progesterone receptor N termini binds SUMO-1 protein to regulate autoinhibition and transrepression. J. Biol. Chem. 277: 33950–33956.

Akimaru H., D. X. Hou, and S. Ishii, 1997 Drosophila CBP is required for dorsal-dependent twist gene expression. Nat. Genet. 17: 211–214.

Al Asafen H., P. U. Bandodkar, S. Carrell-Noel, A. E. Schloop, J. Friedman, et al., 2020 Robustness of the Dorsal morphogen gradient with respect to morphogen dosage. PLoS Comput. Biol. 16: e1007750.

Ambrosi P., J. S. Chahda, H. R. Koslen, H. J. Chiel, and C. M. Mizutani, 2014 Modeling of the dorsal gradient across species reveals interaction between embryo morphology and Toll signaling pathway during evolution. PLoS Comput. Biol. 10: e1003807.

Anderson K. V., and C. Nüsslein-Volhard, 1984 Information for the dorsal–ventral pattern of the Drosophila embryo is stored as maternal mRNA. Nature 311: 223–227.

Anderson K. V., 2000 Toll signaling pathways in the innate immune response. Curr. Opin. Immunol. 12: 13–19.

Anjum S. G., W. Xu, N. Nikkholgh, S. Basu, Y. Nie, et al., 2013 Regulation of Toll signaling and inflammation by β-arrestin and the SUMO protease Ulp1. Genetics 195: 1307–1317.

Araujo H., and E. Bier, 2000 sog and dpp exert opposing maternal functions to modify toll signaling and pattern the dorsoventral axis of the Drosophila embryo. Development 127: 3631–3644.

Banerjee U., J. R. Girard, L. M. Goins, and C. M. Spratford, 2019 Drosophila as a Genetic Model for Hematopoiesis. Genetics 211: 367–417.

Bassett A. R., C. Tibbit, C. P. Ponting, and J.-L. Liu, 2014 Highly Efficient Targeted Mutagenesis of Drosophila with the CRISPR/Cas9 System. Cell Rep. 6: 1178–1179.

Belvin M. P., Y. Jin, and K. V. Anderson, 1995 Cactus protein degradation mediates Drosophila dorsal-ventral signaling. Genes Dev. 9: 783–793.

Belvin M. P., and K. V. Anderson, 1996 A conserved signaling pathway: the Drosophila toll-dorsal pathway. Annu. Rev. Cell Dev. Biol. 12: 393–416.

Bergmann A., D. Stein, R. Geisler, S. Hagenmaier, B. Schmid, et al., 1996 A gradient of cytoplasmic Cactus degradation establishes the nuclear localization gradient of the dorsal morphogen in Drosophila. Mech. Dev. 60: 109–123.

Bettencourt R., H. Asha, C. Dearolf, and Y. T. Ip, 2004 Hemolymph-dependent and - independent responses in Drosophila immune tissue. J. Cell. Biochem. 92: 849–863.

Bhaskar V., S. A. Valentine, and A. J. Courey, 2000 A functional interaction between dorsal and components of the Smt3 conjugation machinery. J. Biol. Chem. 275: 4033–4040.

Bhaskar V., M. Smith, and A. J. Courey, 2002 Conjugation of Smt3 to dorsal may potentiate the Drosophila immune response. Mol. Cell. Biol. 22: 492–504.

Bier E., M. M. Harrison, K. M. O’Connor-Giles, and J. Wildonger, 2018 Advances in Engineering the Fly Genome with the CRISPR-Cas System. Genetics 208: 1–18.

Bies J., J. Markus, and L. Wolff, 2002 Covalent attachment of the SUMO-1 protein to the negative regulatory domain of the c-Myb transcription factor modifies its stability and transactivation capacity. J. Biol. Chem. 277: 8999–9009.

Carrell S. N., M. D. O’Connell, T. Jacobsen, A. E. Pomeroy, S. M. Hayes, et al., 2017 A facilitated diffusion mechanism establishes the Drosophila Dorsal gradient. Development 144: 4450–4461.

Chiu H., B. C. Ring, R. P. Sorrentino, M. Kalamarz, D. Garza, et al., 2005 dUbc9 negatively regulates the Toll-NF-κB pathways in larval hematopoiesis and drosomycin activation in Drosophila. Dev. Biol. 288: 60–72.

Cowden J., and M. Levine, 2003 Ventral dominance governs sequential patterns of gene expression across the dorsal–ventral axis of the neuroectoderm in the Drosophila embryo. Dev. Biol. 262: 335–349.

Drier E. A., L. H. Huang, and R. Steward, 1999 Nuclear import of the Drosophila Rel protein Dorsal is regulated by phosphorylation. Genes Dev. 13: 556–568.

Eloranta J. J., and H. C. Hurst, 2002 Transcription factor AP-2 interacts with the SUMO-conjugating enzyme UBC9 and is sumolated in vivo. J. Biol. Chem. 277: 30798–30804.

Enserink J. M., 2015 Sumo and the cellular stress response. Cell Div. 10: 4.

Ferguson E. L., and K. V. Anderson, 1991 Dorsal-ventral pattern formation in the Drosophila embryo: the role of zygotically active genes, (H. R. Bode, Ed.). Curr. Top. Dev. Biol. 25: 17–43.

Ferrandon D., J.-L. Imler, C. Hetru, and J. A. Hoffmann, 2007 The Drosophila systemic immune response: sensing and signalling during bacterial and fungal infections. Nat. Rev. Immunol. 7: 862–874.

Fossett N., K. Hyman, K. Gajewski, S. H. Orkin, and R. A. Schulz, 2003 Combinatorial interactions of Serpent, Lozenge, and U-shaped regulate crystal cell lineage commitment during Drosophila hematopoiesis. Proc. Natl. Acad. Sci. U. S. A. 100: 11451–11456.

Ganguly A., J. Jiang, and Y. T. Ip, 2005 Drosophila WntD is a target and an inhibitor of the Dorsal/Twist/Snail network in the gastrulating embryo. Development 132: 3419–3429.

Gareau J. R., and C. D. Lima, 2010 The SUMO pathway: emerging mechanisms that shape specificity, conjugation and recognition. Nat. Rev. Mol. Cell Biol. 11: 861–871.

Geisler R., A. Bergmann, Y. Hiromi, and C. Nüsslein-Volhard, 1992 cactus, a gene involved in dorsoventral pattern formation of Drosophila, is related to the IκB gene family of vertebrates. Cell 71: 613–621.

Gillespie S. K., and S. A. Wasserman, 1994 Dorsal, a Drosophila Rel-like protein, is phosphorylated upon activation of the transmembrane protein Toll. Mol. Cell. Biol. 14: 3559–3568.

González-Crespo S., and M. Levine, 1993 Interactions between dorsal and helix-loop-helix proteins initiate the differentiation of the embryonic mesoderm and neuroectoderm in Drosophila. Genes Dev. 7: 1703–1713.

Gordon M. D., M. S. Dionne, D. S. Schneider, and R. Nusse, 2005 WntD is a feedback inhibitor of Dorsal/NF-kappaB in Drosophila development and immunity. Nature 437: 746–749.

Govind S., L. Brennan, and R. Steward, 1993 Homeostatic balance between dorsal and cactus proteins in the Drosophila embryo. Development 117: 135–148.

Govind S., 1999 Control of development and immunity by rel transcription factors in Drosophila. Oncogene 18: 6875–6887.

Govind S., 2008 Innate immunity in Drosophila: Pathogens and pathways. Insect Sci.

Gratz S. J., F. P. Ukken, C. D. Rubinstein, G. Thiede, L. K. Donohue, et al., 2014 Highly specific and efficient CRISPR/Cas9-catalyzed homology-directed repair in Drosophila. Genetics 196: 961–971.

Handu M., B. Kaduskar, R. Ravindranathan, A. Soory, R. Giri, et al., 2015 SUMO-Enriched Proteome for Drosophila Innate Immune Response. G3 5: 2137–2154.

Hay R. T., 2005 SUMO: a history of modification. Mol. Cell 18: 1–12.

Hegde S., A. Soory, B. Kaduskar, and G. S. Ratnaparkhi, 2020 SUMO conjugation regulates immune signalling. Fly 14: 62–79.

Hendriks I. A., and A. C. O. Vertegaal, 2016 A comprehensive compilation of SUMO proteomics. Nat. Rev. Mol. Cell Biol. 17: 581–595.

Hendriks I. A., D. Lyon, C. Young, L. J. Jensen, A. C. O. Vertegaal, et al., 2017 Site-specific mapping of the human SUMO proteome reveals co-modification with phosphorylation. Nat. Struct. Mol. Biol. 24: 325–336.

Hendriks I. A., D. Lyon, D. Su, N. H. Skotte, J. A. Daniel, et al., 2018 Site-specific characterization of endogenous SUMOylation across species and organs. Nat. Commun. 9: 2456.

Hoffmann J. A., 2003 The immune response of Drosophila. Nature 426: 33–38.

Hong J.-W., D. A. Hendrix, D. Papatsenko, and M. S. Levine, 2008 How the Dorsal gradient works: Insights from postgenome technologies. Proc. Natl. Acad. Sci. U. S. A. 105: 20072–20076.

Horner V. L., and M. F. Wolfner, 2008 Transitioning from egg to embryo: triggers and mechanisms of egg activation. Dev. Dyn. 237: 527–544.

Huang L., S. Ohsako, and S. Tanda, 2005 The lesswright mutation activates Rel-related proteins, leading to overproduction of larval hemocytes in Drosophila melanogaster. Dev. Biol. 280: 407–420.

Ip Y. T., R. E. Park, D. Kosman, E. Bier, and M. Levine, 1992 The dorsal gradient morphogen regulates stripes of rhomboid expression in the presumptive neuroectoderm of the Drosophila embryo. Genes Dev. 6: 1728–1739.

Ip Y. T., M. Reach, Y. Engstrom, L. Kadalayil, H. Cai, et al., 1993 Dif, a dorsal-related gene that mediates an immune response in Drosophila. Cell 75: 753–763.

Isoda K., S. Roth, and C. Nüsslein-Volhard, 1992 The functional domains of the Drosophila morphogen dorsal: evidence from the analysis of mutants. Genes Dev. 6: 619–630.

Janeway C. A. Jr, and R. Medzhitov, 2002 Innate immune recognition. Annu. Rev. Immunol. 20: 197–216.

Jiang J., D. Kosman, Y. T. Ip, and M. Levine, 1991 The dorsal morphogen gradient regulates the mesoderm determinant twist in early Drosophila embryos. Genes Dev. 5: 1881–1891.

Johnson E. S., 2004 Protein modification by SUMO. Annu. Rev. Biochem. 73: 355–382.

Karin M., and Y. Ben-Neriah, 2000 Phosphorylation meets ubiquitination: the control of NF-[kappa]B activity. Annu. Rev. Immunol. 18: 621–663.

Kenmoku H., A. Hori, T. Kuraishi, and S. Kurata, 2017 A novel mode of induction of the humoral innate immune response in Drosophila larvae. Dis. Model. Mech. 10: 271–281.

Kondo S., and R. Ueda, 2013 Highly improved gene targeting by germline-specific Cas9 expression in Drosophila. Genetics 195: 715–721.

Kopp E. B., and S. Ghosh, 1995 NF-kB and Rel proteins in innate immunity. Adv. Immunol. 58.

Kosman D., Y. T. Ip, M. Levine, and K. Arora, 1991 Establishment of the mesoderm-neuroectoderm boundary in the Drosophila embryo. Science 254: 118–122.

Kosman D., K. Yazdanbakhsh, and M. Levine, 1992 dorsal-twist interactions establish snail expression in the presumptive mesoderm of the Drosophila embryo. Genes .

Kracklauer M. P., and C. Schmidt, 2003 At the crossroads of SUMO and NF-kappaB. Mol. Cancer 2: 39.

Lanot R., D. Zachary, F. Holder, and M. Meister, 2001 Postembryonic hematopoiesis in Drosophila. Dev. Biol. 230: 243–257.

Laver J. D., A. J. Marsolais, C. A. Smibert, and H. D. Lipshitz, 2015 Regulation and Function of Maternal Gene Products During the Maternal-to-Zygotic Transition in Drosophila. Curr. Top. Dev. Biol. 113: 43–84.

Leidner J., C. Voogdt, R. Niedenthal, P. Möller, U. Marienfeld, et al., 2014 SUMOylation attenuates the transcriptional activity of the NF-κB subunit RelB. J. Cell. Biochem. 115: 1430–1440.

Lemaitre B., M. Meister, S. Govind, P. Georgel, R. Steward, et al., 1995 Functional analysis and regulation of nuclear import of dorsal during the immune response in Drosophila. EMBO J. 14: 536–545.

Lemaitre B., E. Nicolas, L. Michaut, J. M. Reichhart, and J. A. Hoffmann, 1996 The dorsoventral regulatory gene cassette spätzle/Toll/cactus controls the potent antifungal response in Drosophila adults. Cell 86: 973–983.

Li R., X. Yao, H. Zhou, P. Jin, and F. Ma, 2021 The Drosophila miR-959–962 Cluster Members Repress Toll Signaling to Regulate Antibacterial Defense during Bacterial Infection. Int. J. Mol. Sci. 22: 886.

Liberman L. M., G. T. Reeves, and A. Stathopoulos, 2009 Quantitative imaging of the Dorsal nuclear gradient reveals limitations to threshold-dependent patterning in Drosophila. Proc. Natl. Acad. Sci. U. S. A. 106: 22317–22322.

Liu Z. P., R. L. Galindo, and S. A. Wasserman, 1997 A role for CKII phosphorylation of the cactus PEST domain in dorsoventral patterning of the Drosophila embryo. Genes Dev. 11: 3413–3422.

Liu Y., R. Bridges, A. Wortham, and M. Kulesz-Martin, 2012 NF-κB repression by PIAS3 mediated RelA SUMOylation. PLoS One 7: e37636.

Mabb A. M., and S. Miyamoto, 2007 SUMO and NF-kappaB ties. Cell. Mol. Life Sci. 64: 1979–1996.

Maggert K., M. Levine, and M. Frasch, 1995 The somatic-visceral subdivision of the embryonic mesoderm is initiated by dorsal gradient thresholds in Drosophila. Development 121: 2107–2116.

Manfruelli P., J. M. Reichhart, R. Steward, J. A. Hoffmann, and B. Lemaitre, 1999 A mosaic analysis in Drosophila fat body cells of the control of antimicrobial peptide genes by the Rel proteins Dorsal and DIF. EMBO J. 18: 3380–3391.

Markstein M., P. Markstein, V. Markstein, and M. S. Levine, 2002 Genome-wide analysis of clustered Dorsal binding sites identifies putative target genes in the Drosophila embryo. Proc. Natl. Acad. Sci. U. S. A. 99: 763–768.

Matova N., and K. V. Anderson, 2006 Rel/NF-κB double mutants reveal that cellular immunity is central to Drosophila host defense. Proceedings of the National.

Matova N., and K. V. Anderson, 2010 Drosophila Rel proteins are central regulators of a robust, multi-organ immune network. J. Cell Sci. 123: 627–633.

Medzhitov R., P. Preston-Hurlburt, and C. A. Janeway Jr, 1997 A human homologue of the Drosophila Toll protein signals activation of adaptive immunity. Nature 388: 394–397.

Meister M., 2004 Blood cells of Drosophila: cell lineages and role in host defence. Curr. Opin. Immunol. 16: 10–15.

Meng X., B. S. Khanuja, and Y. T. Ip, 1999 Toll receptor-mediated Drosophila immune response requires Dif, an NF-kappaB factor. Genes Dev. 13: 792–797.

Morisato D., and K. V. Anderson, 1995 Signaling pathways that establish the dorsal-ventral pattern of the Drosophila embryo. Annu. Rev. Genet. 29: 371–399.

Moussian B., and S. Roth, 2005 Dorsoventral Axis Formation in the Drosophila Embryo—Shaping and Transducing a Morphogen Gradient. Curr. Biol. 15: R887–R899.

Müller S., M. Berger, F. Lehembre, and J. S. Seeler, 2000 c-Jun and p53 activity is modulated by SUMO-1 modification. Journal of Biological.

Nicolas E., J. M. Reichhart, J. A. Hoffmann, and B. Lemaitre, 1998 In vivo regulation of the IκB homologue cactus during the immune response of Drosophila. J. Biol. Chem. 273: 10463–10469.

Nie M., Y. Xie, J. A. Loo, and A. J. Courey, 2009 Genetic and proteomic evidence for roles of Drosophila SUMO in cell cycle control, Ras signaling, and early pattern formation. PLoS One 4: e5905.

Nishida T., and H. Yasuda, 2002 PIAS1 and PIASxα Function as SUMO-E3 Ligases toward Androgen Receptor and Repress Androgen Receptor-dependent Transcription *. J. Biol. Chem. 277: 41311–41317.

Nüsslein-Volhard C., 1979 Maternal effect mutations that alter the spatial coordinates of the embryo of Drosophila melanogaster. Determinants of spatial organization 28.

Nüsslein-Volhard C., M. Lohs-Schardin, K. Sander, and C. Cremer, 1980 A dorso-ventral shift of embryonic primordia in a new maternal-effect mutant of Drosophila. Nature 283: 474–476.

Ouyang J., and G. Gill, 2009 SUMO engages multiple corepressors to regulate chromatin structure and transcription. Epigenetics 4: 440–444.

Paddibhatla I., M. J. Lee, M. E. Kalamarz, R. Ferrarese, and S. Govind, 2010 Role for sumoylation in systemic inflammation and immune homeostasis in Drosophila larvae. PLoS Pathog. 6: e1001234.

Papatsenko D., and M. Levine, 2005 Quantitative analysis of binding motifs mediating diverse spatial readouts of the Dorsal gradient in the Drosophila embryo. Proc. Natl. Acad. Sci. U. S. A. 102: 4966–4971.

Papatsenko D., and M. S. Levine, 2008 How the Dorsal gradient works: insights from postgenome technologies. Proceedings of the.

Petersen U. M., L. Kadalayil, K. P. Rehorn, D. K. Hoshizaki, R. Reuter, et al., 1999 Serpent regulates Drosophila immunity genes in the larval fat body through an essential GATA motif. EMBO J. 18: 4013–4022.

Pirone L., W. Xolalpa, J. O. Sigurðsson, J. Ramirez, C. Pérez, et al., 2017 A comprehensive platform for the analysis of ubiquitin-like protein modifications using in vivo biotinylation. Sci. Rep. 7: 40756.

Poukka H., U. Karvonen, O. A. Janne, and J. J. Palvimo, 2000 Covalent modification of the androgen receptor by small ubiquitin-like modifier 1 (SUMO-1). Proc. Natl. Acad. Sci. U. S. A. 97: 14145–14150.

Qiu P., P. C. Pan, and S. Govind, 1998 A role for the Drosophila Toll/Cactus pathway in larval hematopoiesis. Development 125: 1909–1920.

Raman N., A. Nayak, and S. Muller, 2013 The SUMO system: a master organizer of nuclear protein assemblies. Chromosoma 122: 475–485.

Ratnaparkhi G. S., H. A. Duong, and A. J. Courey, 2008 Dorsal interacting protein 3 potentiates activation by Drosophila Rel homology domain proteins. Dev. Comp. Immunol. 32: 1290–1300.

Ratnaparkhi G. S., and A. J. Courey, 2015 Chapter 8 - Signaling Cascades, Gradients, and Gene Networks in Dorsal/Ventral Patterning, pp. 131–151 in Principles of Developmental Genetics (Second Edition), edited by Moody S. A. Academic Press, Oxford.

Ray R. P., K. Arora, C. Nüsslein-Volhard, and W. M. Gelbart, 1991 The control of cell fate along the dorsal-ventral axis of the Drosophila embryo. Development 113: 35–54.

Reeves G. T., and A. Stathopoulos, 2009 Graded dorsal and differential gene regulation in the Drosophila embryo. Cold Spring Harb. Perspect. Biol. 1: a000836.

Reeves G. T., N. Trisnadi, T. V. Truong, M. Nahmad, S. Katz, et al., 2012 Dorsal-ventral gene expression in the Drosophila embryo reflects the dynamics and precision of the dorsal nuclear gradient. Dev. Cell 22: 544–557.

Reichhart J. M., P. Georgel, M. Meister, B. Lemaitre, C. Kappler, et al., 1993 Expression and nuclear translocation of the rel/NF-kappa B-related morphogen dorsal during the immune response of Drosophila. C. R. Acad. Sci. III 316: 1218–1224.

Rizki M. T., and R. M. Rizki, 1959 Functional significance of the crystal cells in the larva of Drosophila melanogaster. J. Biophys. Biochem. Cytol. 5: 235–240.

Rosonina E., A. Akhter, Y. Dou, J. Babu, and V. S. Sri Theivakadadcham, 2017 Regulation of transcription factors by sumoylation. Transcription 8: 220–231.

Rosonina E., 2019 A conserved role for transcription factor sumoylation in binding-site selection. Curr. Genet. 65: 1307–1312.

Ross S., J. L. Best, L. I. Zon, and G. Gill, 2002 SUMO-1 modification represses Sp3 transcriptional activation and modulates its subnuclear localization. Mol. Cell 10: 831–842.

Roth S., D. Stein, and C. Nüsslein-Volhard, 1989 A gradient of nuclear localization of the dorsal protein determines dorsoventral pattern in the Drosophila embryo. Cell 59: 1189–1202.

Roth S., Y. Hiromi, D. Godt, and C. Nüsslein-Volhard, 1991 cactus, a maternal gene required for proper formation of the dorsoventral morphogen gradient in Drosophila embryos. Development 112: 371–388.

Rusch J., and M. Levine, 1996 Threshold responses to the dorsal regulatory gradient and the subdivision of primary tissue territories in the Drosophila embryo. Curr. Opin. Genet. Dev. 6: 416–423.

Rushlow C. A., K. Han, J. L. Manley, and M. Levine, 1989 The graded distribution of the dorsal morphogen is initiated by selective nuclear transport in Drosophila. Cell 59: 1165–1177.

Rutschmann S., A. Kilinc, and D. Ferrandon, 2002 Cutting edge: the toll pathway is required for resistance to gram-positive bacterial infections in Drosophila. J. Immunol. 168: 1542–1546.

Ryu H.-Y., S. H. Ahn, and M. Hochstrasser, 2020 SUMO and cellular adaptive mechanisms. Exp. Mol. Med. 52: 931–939.

Santamaria P., and C. Nüsslein-Volhard, 1983 Partial rescue of dorsal , a maternal effect mutation affecting the dorso-ventral pattern of the Drosophila embryo, by the injection of wild-type cytoplasm. EMBO J. 2: 1695–1699.

Sapetschnig A., G. Rischitor, H. Braun, A. Doll, M. Schergaut, et al., 2002 Transcription factor Sp3 is silenced through SUMO modification by PIAS1. EMBO J. 21: 5206–5215.

Schmid M. R., I. Anderl, L. Vesala, L.-M. Vanha-aho, X.-J. Deng, et al., 2014 Control of Drosophila blood cell activation via Toll signaling in the fat body. PLoS One 9: e102568.

Senger K., G. W. Armstrong, W. J. Rowell, J. M. Kwan, M. Markstein, et al., 2004 Immunity Regulatory DNAs Share Common Organizational Features in Drosophila. Mol. Cell 13: 19–32.

Simpson P., 1983 Maternal-Zygotic Gene Interactions during Formation of the Dorsoventral Pattern in Drosophila Embryos. Genetics 105: 615–632.

Sorrentino R. P., J. P. Melk, and S. Govind, 2004 Genetic analysis of contributions of dorsal group and JAK-Stat92E pathway genes to larval hemocyte concentration and the egg encapsulation response in Drosophila. Genetics 166: 1343–1356.

St Johnston D., and C. Nüsslein-Volhard, 1992 The origin of pattern and polarity in the Drosophila embryo. Cell 68: 201–219.

Stathopoulos A., and M. Levine, 2002 Dorsal gradient networks in the Drosophila embryo. Dev. Biol. 246: 57–67.

Stathopoulos A., M. Van Drenth, A. Erives, M. Markstein, and M. Levine, 2002 Whole-genome analysis of dorsal-ventral patterning in the Drosophila embryo. Cell 111: 687–701.

Stein D. S., and L. M. Stevens, 2014 Maternal control of the Drosophila dorsal-ventral body axis. Wiley Interdiscip. Rev. Dev. Biol. 3: 301–330.

Steward R., 1987 Dorsal, an embryonic polarity gene in Drosophila, is homologous to the vertebrate proto-oncogene, c-rel. Science 238: 692–694.

Steward R., S. B. Zusman, L. H. Huang, and P. Schedl, 1988 The dorsal protein is distributed in a gradient in early Drosophila embryos. Cell 55: 487–495.

Steward R., and S. Govind, 1993 Dorsal-ventral polarity in the Drosophila embryo. Curr. Opin. Genet. Dev. 3: 556–561.

Tadros W., and H. D. Lipshitz, 2009 The maternal-to-zygotic transition: a play in two acts. Development 136: 3033–3042.

Tang R., W. Huang, J. Guan, Q. Liu, B. T. Beerntsen, et al., 2021 Drosophila H2Av negatively regulates the activity of the IMD pathway via facilitating Relish SUMOylation. PLoS Genet. 17: e1009718.

Tautz D., and C. Pfeifle, 1989 A non-radioactive in situ hybridization method for the localization of specific RNAs in Drosophila embryos reveals translational control of the segmentation gene hunchback. Chromosoma 98: 81–85.

Tay S., J. J. Hughey, T. K. Lee, T. Lipniacki, S. R. Quake, et al., 2010 Single-cell NF-kappaB dynamics reveal digital activation and analogue information processing. Nature 466: 267–271.

Tempé D., M. Piechaczyk, and G. Bossis, 2008 SUMO under stress. Biochem. Soc. Trans. 36: 874–878.

Tian S., H. Poukka, J. J. Palvimo, and O. A. Jänne, 2002 Small ubiquitin-related modifier-1 (SUMO-1) modification of the glucocorticoid receptor. Biochem. J 367: 907–911.

Valanne S., J. H. Wang, and M. Rämet, 2011 The Drosophila toll signaling pathway. The Journal of Immunology.

Varejão N., J. Lascorz, Y. Li, and D. Reverter, 2020 Molecular mechanisms in SUMO conjugation. Biochem. Soc. Trans. 48: 123–135.

Verger A., J. Perdomo, and M. Crossley, 2003 Modification with SUMO. A role in transcriptional regulation. EMBO Rep. 4: 137–142.

Vlisidou I., and W. Wood, 2015 Drosophila blood cells and their role in immune responses. FEBS J. 282: 1368–1382.

Waddington C. H., 1959 Canalization of Development and Genetic Assimilation of Acquired Characters. Nature 183: 1654–1655.

Whalen A. M., and R. Steward, 1993 Dissociation of the dorsal-cactus complex and phosphorylation of the dorsal protein correlate with the nuclear localization of dorsal. J. Cell Biol. 123: 523–534.

Wieschaus E., and C. Nüsslein-Volhard, 2016 The Heidelberg Screen for Pattern Mutants of Drosophila: A Personal Account. Annu. Rev. Cell Dev. Biol. 32: 1–46.

Williams M. J., 2007 Drosophila hemopoiesis and cellular immunity. J. Immunol. 178: 4711–4716.

Wotton D., L. F. Pemberton, and J. Merrill-Schools, 2017 SUMO and Chromatin Remodeling. Adv. Exp. Med. Biol. 963: 35–50.

Wykoff D. D., and E. K. O’Shea, 2005 Identification of Sumoylated Proteins by Systematic Immunoprecipitation of the Budding Yeast Proteome *. Mol. Cell. Proteomics 4: 73–83.

Zeitlinger J., R. P. Zinzen, A. Stark, and M. Kellis, 2007 Whole-genome ChIP–chip analysis of Dorsal, Twist, and Snail suggests integration of diverse patterning processes in the Drosophila embryo. Genes .

Zhang G., and S. Ghosh, 2001 Toll-like receptor–mediated NF-κB activation: a phylogenetically conserved paradigm in innate immunity. J. Clin. Invest. 107: 13–19.

Zhou J., J. Zwicker, P. Szymanski, M. Levine, and R. Tjian, 1998 TAFII mutations disrupt Dorsal activation in the Drosophila embryo. Proc. Natl. Acad. Sci. U. S. A. 95: 13483–13488.

Zhou R., N. Silverman, M. Hong, D. S. Liao, Y. Chung, et al., 2005 The role of ubiquitination in Drosophila innate immunity. J. Biol. Chem. 280: 34048–34055.

