## Supplementary figures for "SUMOylation of Dorsal attenuates Toll/NF-κB signalling"

Fig. S1

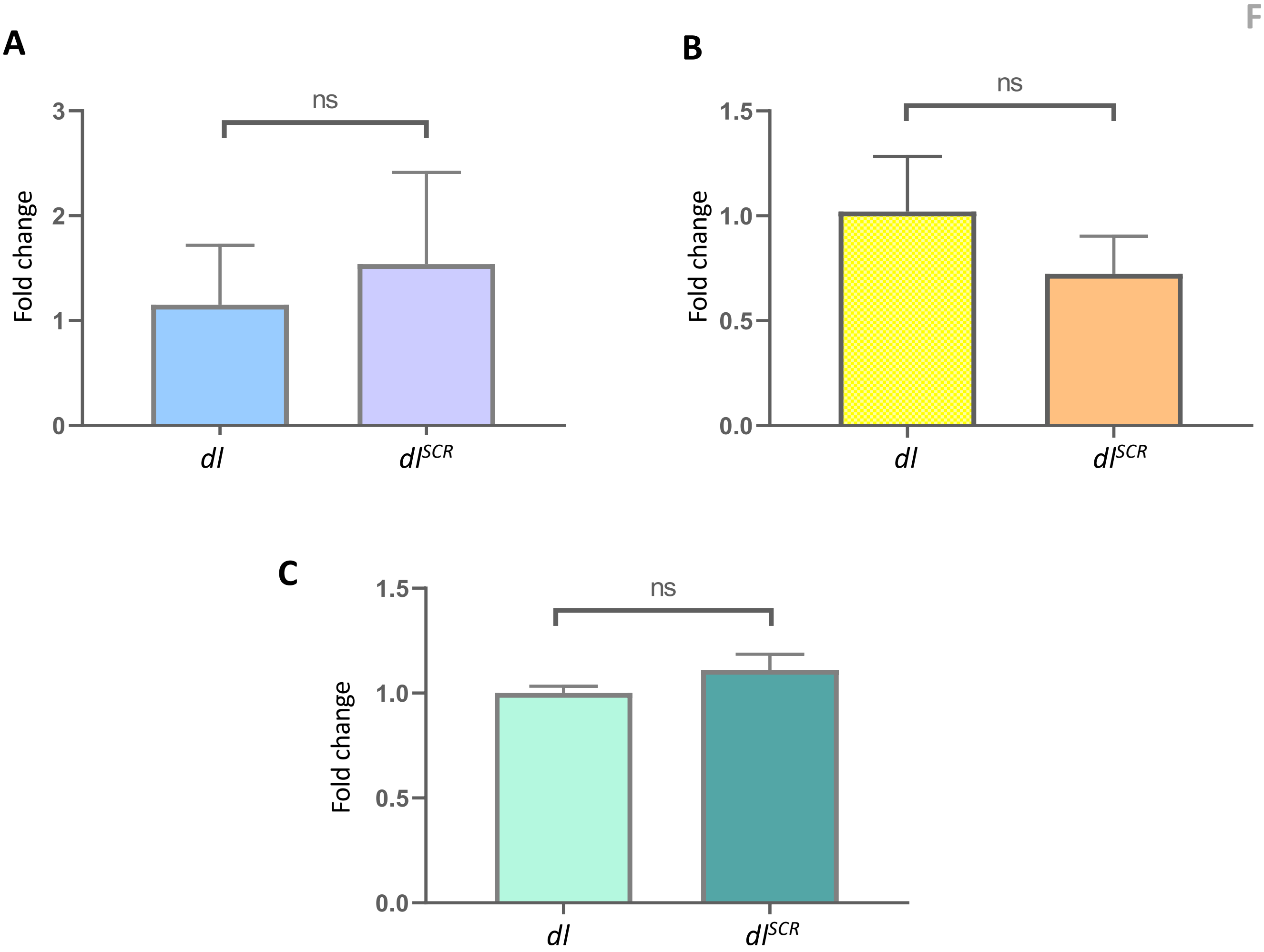


**Fig. S1.** *dl* transcript levels assayed by qRT-PCR across different stages of the fly life cycle- embryo (A), larva (B) and adult (D) for the genotypes *dl* and *dl*$^{SCR}$.

Fig. S2

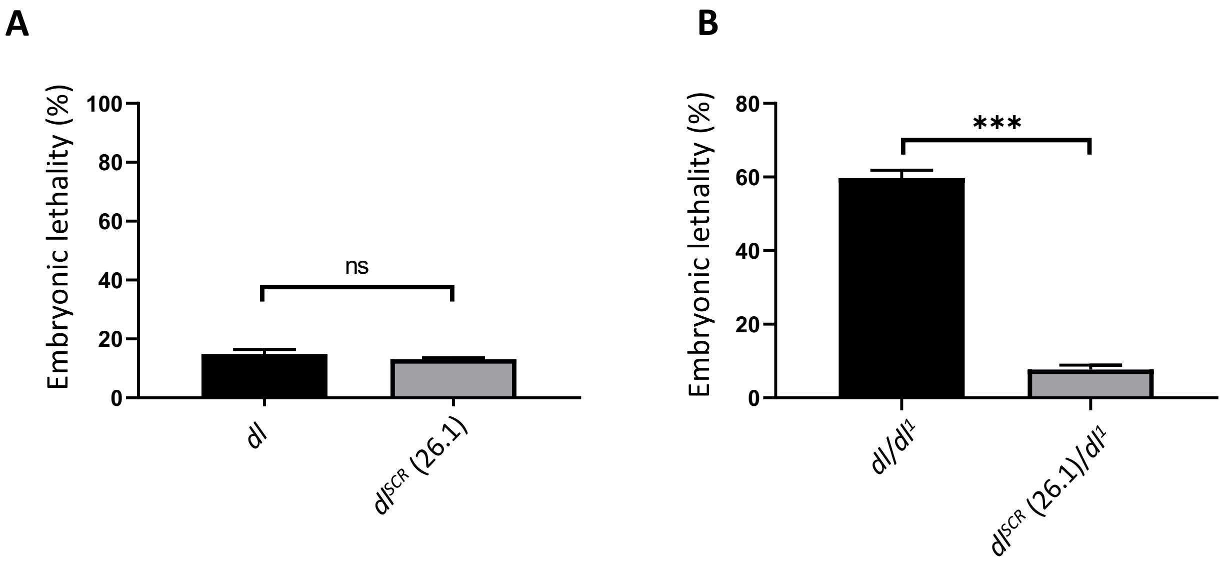


**Fig. S2.** Embryonic lethality is plotted for the indicated genotypes, at 29 °C. N=3, mean ± SEM, Unpaired t-test, (ns) *P* >0.05, (***) *P* < 0.001.

A

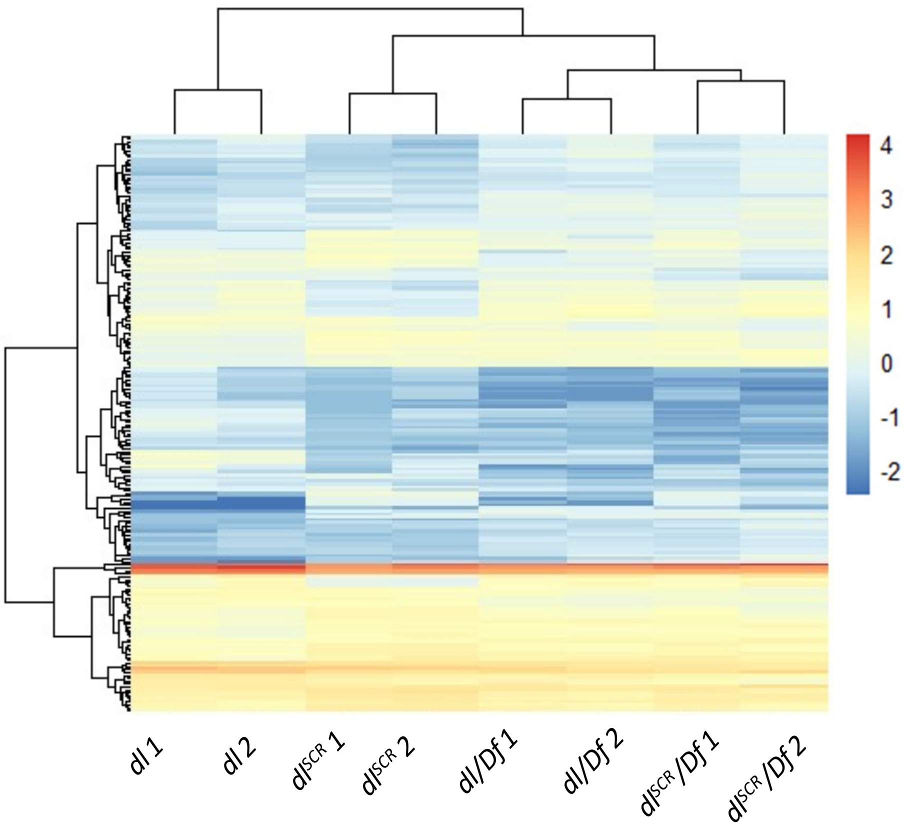


C

| Gene | log2 fold change | Gene network |
|---|---|---|
| *zen* | 1.64 | D-V |
| *kni* | 1.34 | A-P |
| *twi* | 1.33 | D-V |
| *rho* | 1.32 | D-V |
| *sna* | 1.08 | D-V |
| *ftz* | 1.06 | A-P |
| *oc* | 1.04 | A-P |
| *Egfr* | 0.99 | EGF |
| *tll* | 0.985 | A-P |
| *opa* | 0.982 | A-P |
| *tld* | 0.92 | D-V |
| *run* | 0.91 | D-V |
| *ths* | 0.86 | D-V |
| *hkb* | 0.85 | A-P |

B

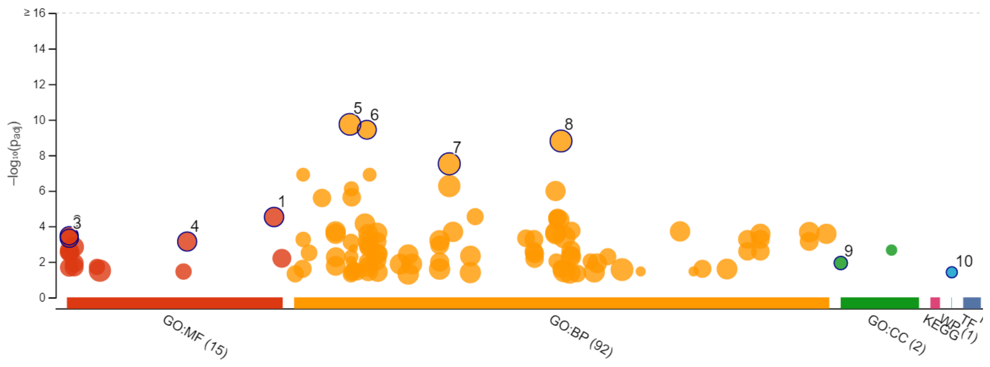


**GO term**

1. transcription regulator activity
2. RNA polymerase II cis-regulatory region sequence-specific DNA binding
3. cis-regulatory region sequence-specific DNA binding
4. sequence-specific DNA binding
5. multicellular organism development
6. embryo development
7. developmental process
8. anatomical structure development
9. histone deacetylase complex
10. Hedgehog Signaling Pathway

**Fig. S3**. (A) Heat map of differentially expressed genes across *dl* , *dl*$^{SCR}$, *dl/Df, dl*$^{SCR}$*/Df are* represented, in duplicates. Scaled, LogCPM values are plotted. (B) denotes significantly enriched gene ontology (GO) terms, for genes in (A), categorized by molecular function (MF), biological process (BP), cellular component (CC) and pathway (KEGG). A subset of differentially regulated genes compared across *dl/Df* and *dl*$^{SCR}$*/Df* with known binding sites for DL is listed in (C), along with the corresponding log2 fold-change values. The genes are broadly classified according to the gene regulatory network they are a part of in the early embryo: D-V (Dorso-ventral patterning; in grey), A-P (Antero-posterior patterning; in yellow) and the EGF signaling pathway (in orange).

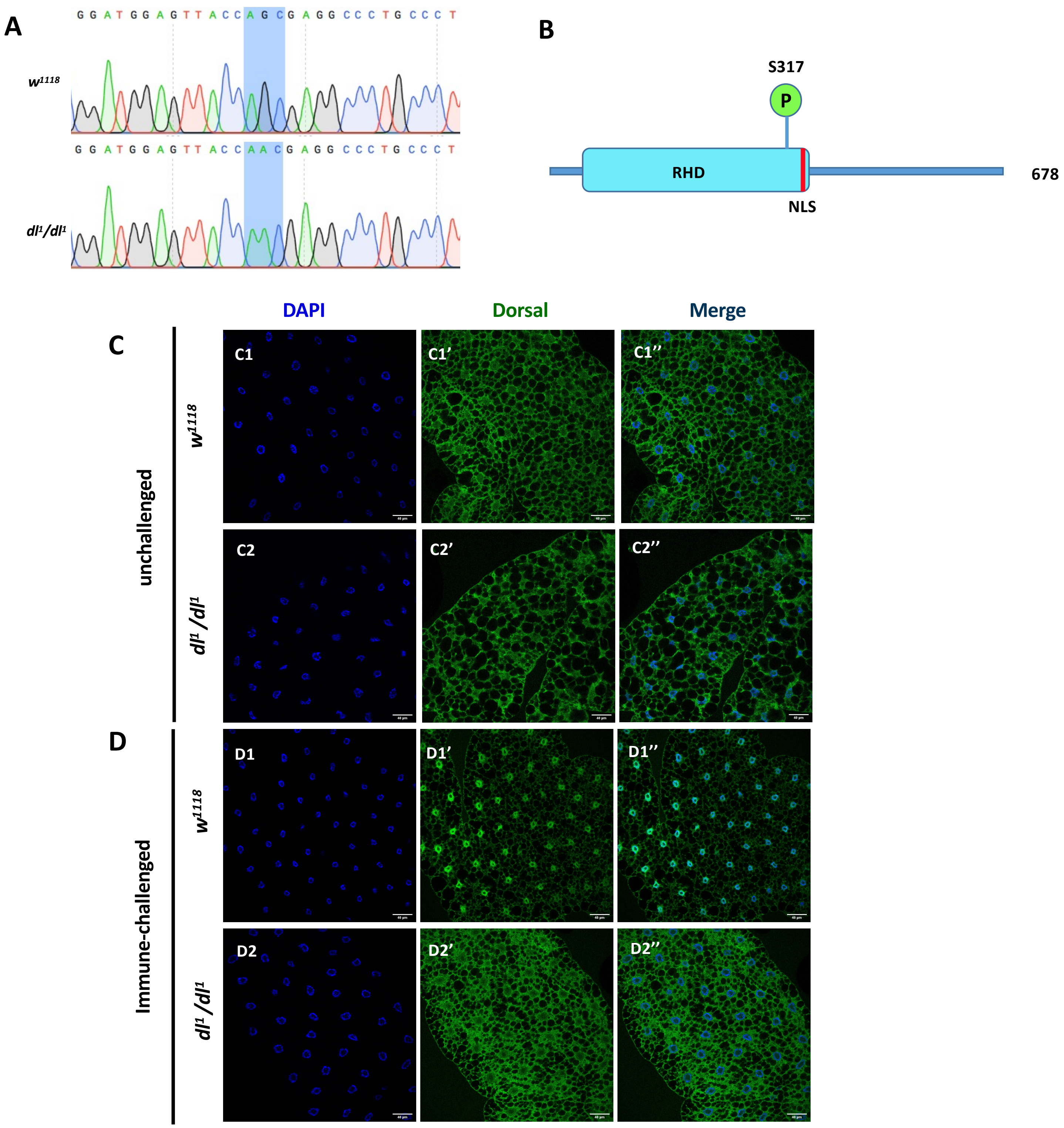


**Fig. S4. DL¹ is resistant to Toll signalling.** The CDS of *dl¹/dl¹* was sequenced and found to harbour a mutation in a critical S317 residue in the rel-homology domain, subject to a phospho-modification (A). The presence of the S317N mutation is highlighted in blue, in (B), along with a reference sequence for *w¹¹¹⁸*. In the unchallenged condition, immunostaining with a DL antibody reveals a pre-dominantly cytoplasmic distribution of DL in *dl¹/dl¹* and *w¹¹¹⁸* (C'-C'') in the fat body. Septic injury with *S. saprophyticus* triggers the nuclear migration of DL in the wild-type animal (D1-D1''), while DL1 is retained in the cytoplasm (D2-D2''). Nuclei are marked with DAPI. N=3, n=5

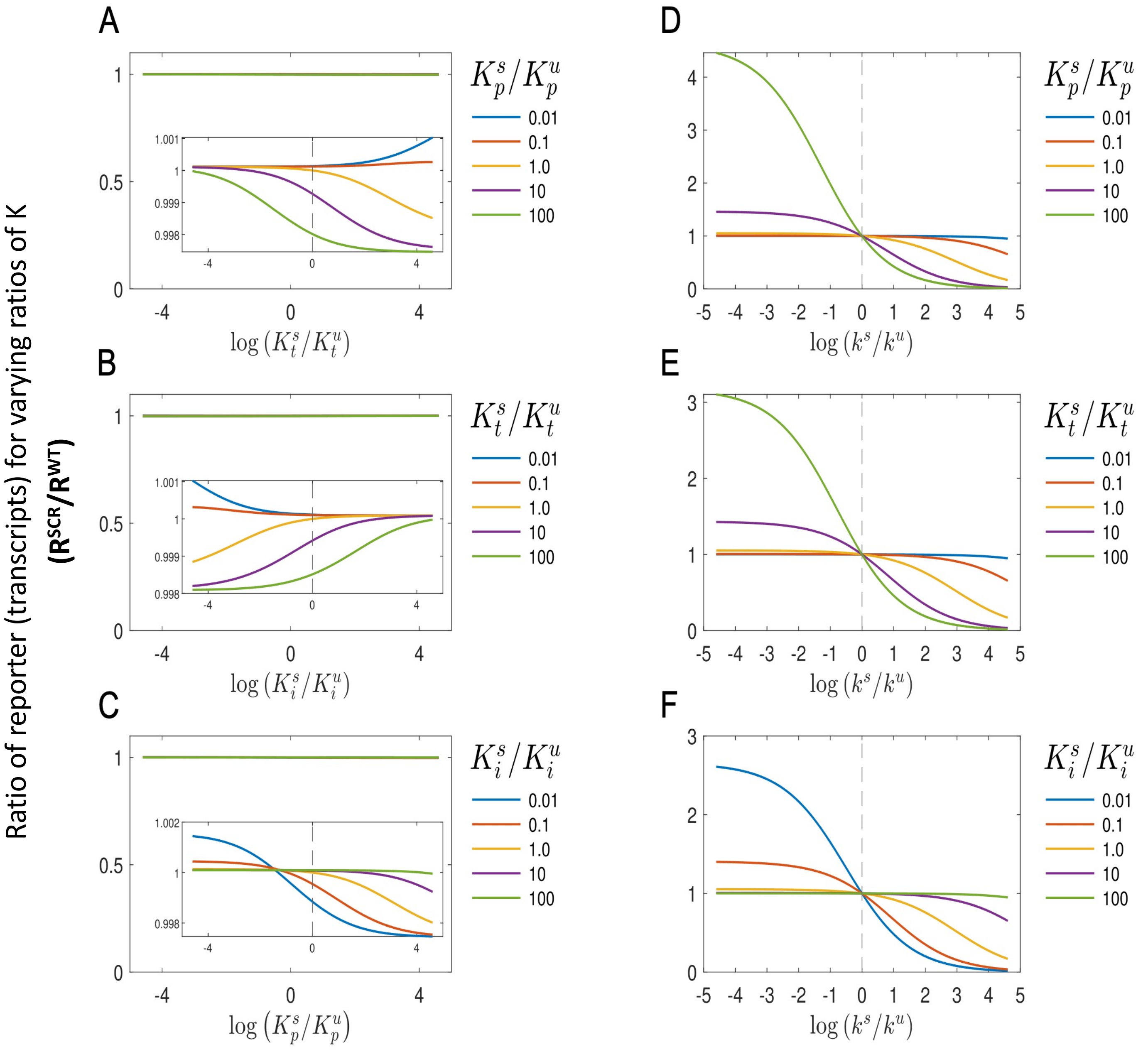


**Fig. S5. Simulating the effect of DL SUMOylation**: The reporter expression levels for the $DL^{SCR}$ is unchanged from the corresponding WT level when the effect of SUMOylation is on Cactus-inhibition strength (A), nuclear-cytoplasmic partition coefficient (B) or promoter binding affinity (C). However when the specific transcription rate is lower($k^s/k^u<1$) for the $DL^S$ relative to $DL^U$, the mutant shows higher relative expression levels, which are modulated by changes in the promoter binding (D), partitioning (E) or Cact binding (F).

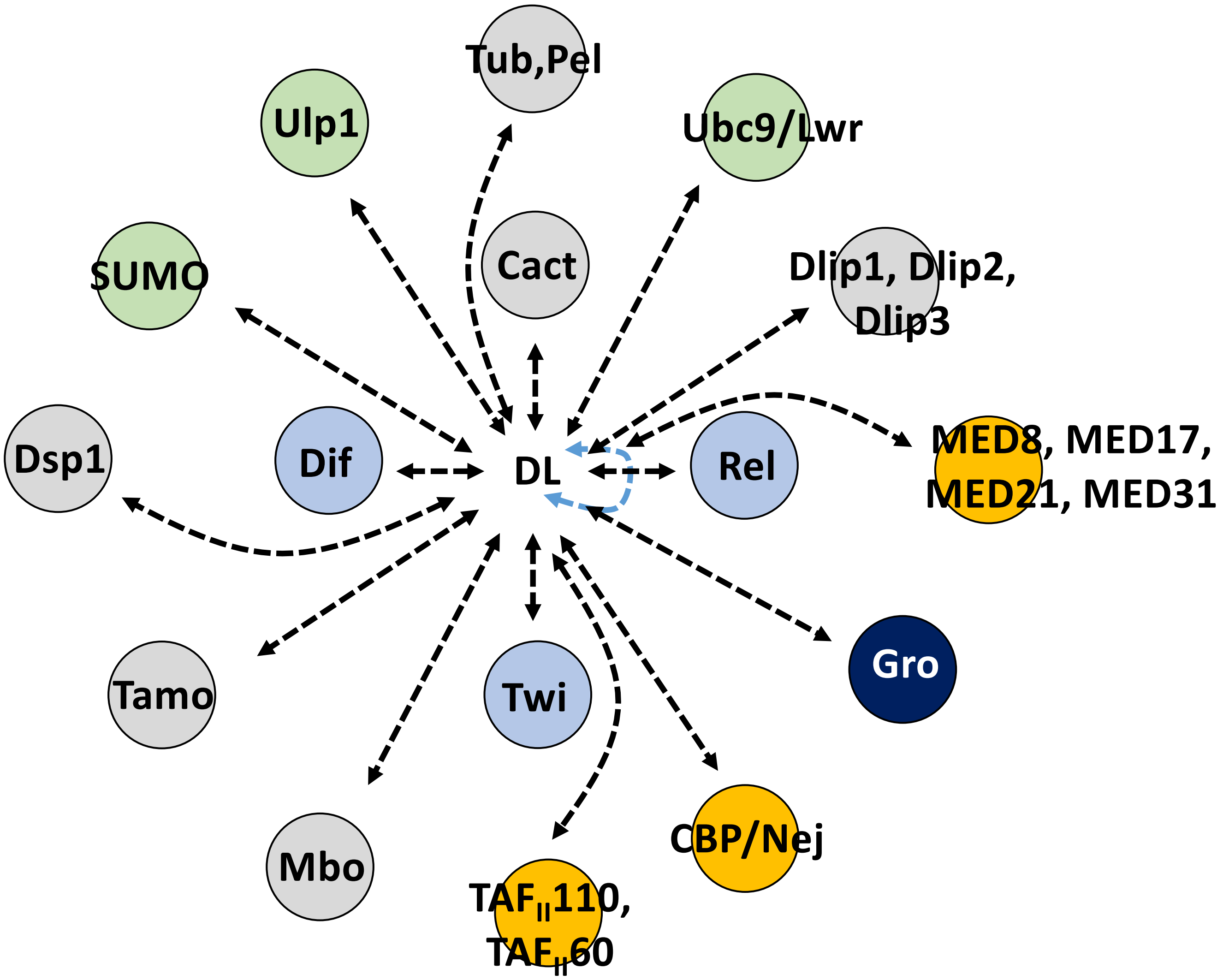


**Fig. S6. Physical interactors of DL.** A list of DL interactors based on data collated in Flybase. The SUMO conjugation and de-conjugation machinery (green circles) interacts with DL, as does the machinery involved in transcriptional activation (orange circles). SUMOylation of DL may perturb the interaction of DL with anyone or all of these partner proteins and in turn may influence DL mediated activation or repression.
